# Learning steers the ontogeny of an efficient hunting sequence in zebrafish larvae

**DOI:** 10.1101/2019.12.19.883157

**Authors:** Konstantinos Lagogiannis, Giovanni Diana, Martin P Meyer

## Abstract

The success of goal-directed behaviours relies on the coordinated execution of a sequence of component actions. In young animals, such sequences may be poorly coordinated, but with age and experience, behaviour progressively adapts to efficiently exploit the animal’s ecological niche. How experience impinges on the developing neural circuits of behaviour is an open question. As a model system, larval zebrafish (*Danio rerio*) hold enormous potential for studying both the development of behaviour and the underlying circuits, but no relevant experience-dependent learning paradigm has yet been characterized. To address this, we have conducted a detailed study of the effects of experience on the ontogeny of hunting behaviour in larval zebrafish. We report that larvae with prior prey experience consume considerably more prey than naive larvae. This is mainly due to increased capture success that is also accompanied by a modest increase in hunt rate. We identified two components of the hunting sequence that are jointly modified by experience. At the onset of the hunting sequence, the orientation strategy of the turn towards prey is modified such that experienced larvae undershoot prey azimuth. Near the end of the hunt sequence, we find that experienced larvae are more likely to employ high-speed capture swims initiated from longer distances to prey. Combined, these modified turn and capture manoeuvrers can be used to predict the probability of capture success and suggest that their development provides advantages specific to larvae feeding on live-prey. Our findings establish an ethologically relevant paradigm in zebrafish for studying how the brain is shaped by experience to drive the ontogeny of efficient behaviour.

## 1 Introduction

The study of animal ethology has demonstrated an instrumental role of early experience in permanently embedding specific information about the environment in the developing animal, and in extending and enhancing behavioural repertoires [4, 47].

An example of a dynamic behaviour that is found to benefit from learning by experience is predation as, across diverse species, it involves predicting and responding to the behaviour of another animal. In altricial species, which require an extended period of parental care, young animals learn to hunt from direct experience, and from mimicking the behaviour of parents or other conspecifics [14]. Even in precocial species, in which hunting behaviour is developed prenatally, hunting can nevertheless be modified by experience. For example, with experience, hatchling snakes improve in their ability to capture prey [35] and orb-web spiders build more effective webs [22]. In fish, the early fine-tuning of development to the available prey types can be critical for survival in the wild. Studies of hatchery-raised fish have found that a lack of prior exposure to prey, before being released into the wild, can strongly reduce their chances of surviving to adulthood [see 11, 12] [7, 13, 17, 37]. It is unlikely that exposure to prey simply facilitates certain aspects of behavioural ontogeny that would have developed with time [4], because in at least some fish species, the detection, handling and capturing of prey is enhanced only for the particular type of prey they have experienced [13, 16, 37]. Such evidence suggests that the ontogeny of hunting behaviour relies on relevant early experience to fully develop [11, 53]. Learning by experience could modify both the perceptual and the kinematic components involved in prey detection, pursuit and capture. However, which aspects of hunting behaviour are malleable by experience and how these contribute to increasing hunting performance remains largely unknown.

The larval zebrafish is an ideal vertebrate model for the detailed study of behaviour and its associated circuits. They have a diverse behavioural repertoire that can be precisely measured [31, 41], and neural activity can be recorded non-invasively and at single-neuron resolution throughout the whole brain [1, 29, 40, 45, 57]. Larval hunting appears as a distinct behavioural mode that is composed of several component actions chained together into a sequence [9, 33, 51]. While significant progress has been made in characterising the kinematics of larval zebrafish hunting in detail [6, 9, 43, 51], and in describing the circuits and cell types involved in this behaviour [3, 5, 23, 38, 39, 46, 49, 50], it is not known whether experience affects the ontogeny and effectiveness of their hunting sequences. To address this, we used high speed imaging and behavioural tracking of freely-swimming zebrafish larvae to compare hunting behaviour and performance between larvae that have been reared with live prey and those that have not.

Consistent with previous reports, we find that hunting episodes begin with a convergent saccade and a turn, which accounts for a large proportion of the orienting response towards prey. The convergent saccade is maintained throughout the hunting sequence during which larvae home in on the target using a series of temporally discrete swim bouts, which successively minimize the distance to prey, up to a point where they can perform a final capture manoeuvre. We found that experience resulted in a modest increase in the probability of initiating hunting behaviour, suggesting that the ability to detect prey or the motivation to hunt may be influenced by prior experience. However, the main effect of experience was a marked increase in the probability of a hunt sequence resulting in successful capture. Detailed examination of successful hunting sequences revealed that experienced larvae were kinematically distinct in at least two aspects. Firstly, at the onset of hunting we found that they make an initial turn-to-prey that undershoots prey azimuth. Paradoxically, inexperienced larvae display initial turns that do not undershoot, but rather align them more with prey azimuth. Secondly, we find that experienced larvae are more likely to employ high-speed capture swims that also tend to be initiated at a longer distance from prey. We show that in experienced larvae capture speed is coordinated with distance to prey, and is also combined with undershooting in their turn-to-prey behaviour. We then identify that these coordinated behaviours contribute to the capture success of hunt-sequences, and can be used to predict larval hunting efficiency.

## 2 Results

Zebrafish larvae were reared in one of three different feeding regimes: a live-fed (LF) group, which received Rotifers, a group that was not-fed (NF), and a third group that received dry growth food (DF). After two days of feeding, the effects of hunting experience in the LF group were examined. The DF and NF groups served as controls for behavioural effects that can be attributed to nutritional state (see Section 4.A). A graphical timeline of our experimental protocol is shown on Figure 1. Consistent with previous observations we did not observe a decrease in survival rates of larvae in the NF group [25]. However a small, but statistically significant, effect on growth was determined by measuring mean body lengths at 7dpf (NF=4.18mm,DF=4.26mm,LF=4.34mm see ***Figure S13***) [42].

**Figure 1.**
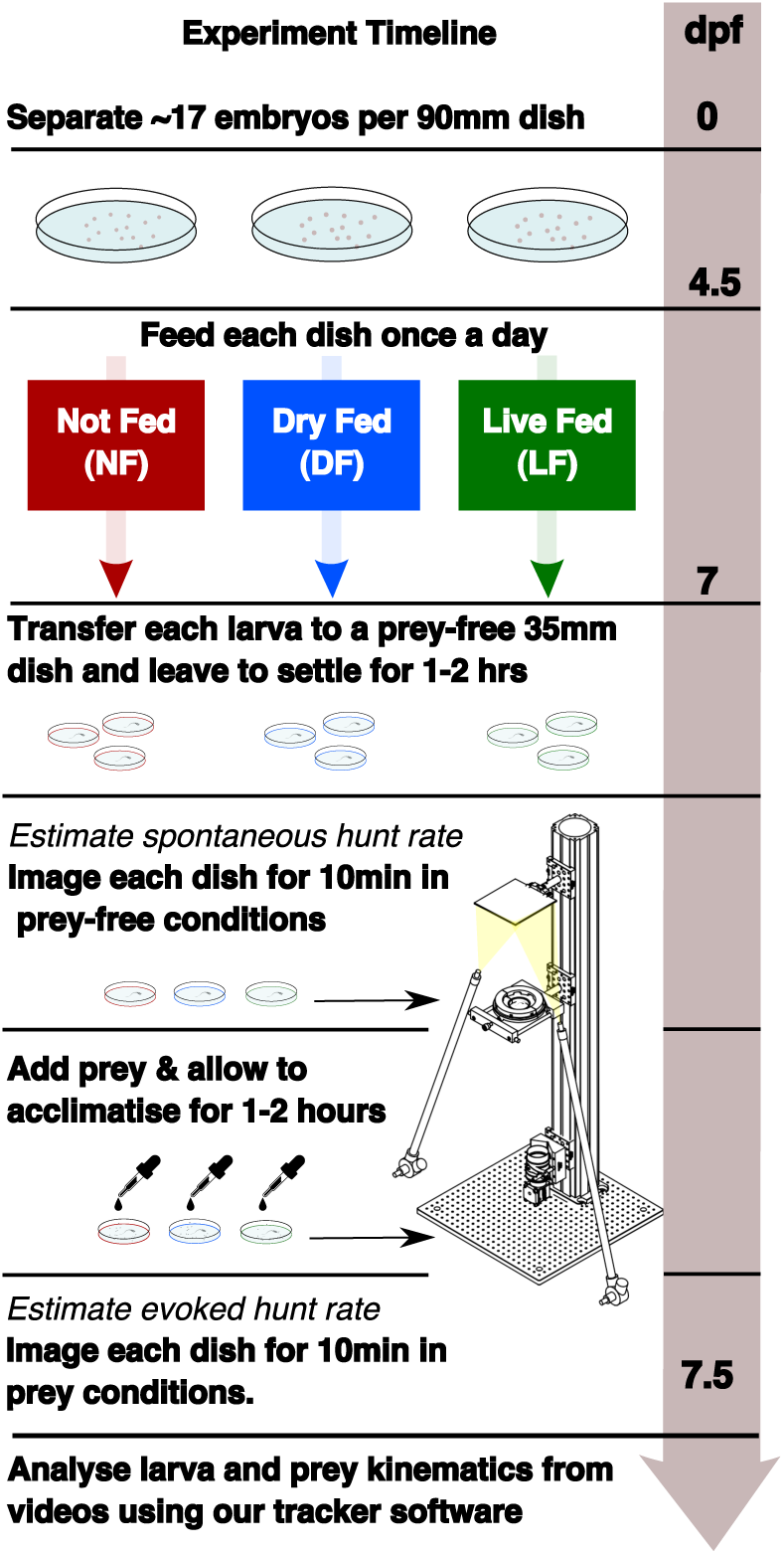
Experimental timeline showing the different feeding regimes during rearing and the recording of spontaneous and evoked hunt behaviour. Embryos are separated in 3 dishes, with differential feeding initiated at 4.5dpf. Each group receives a feed once per day (see Section 4.A). Behavioural recording is performed at 7dpf. Spontaneous eye-vergence events are measured by recording individual larvae for 10min in the absence of prey. Live prey (≈ 30 rotifers) are then added. Acclimatization in prey conditions for a period of ≈ 1-2hr is followed by recording of larvae in the presence of live prey. Live prey is topped up to initial level prior to recording.

Each larva was left to settle for approximately 1-2hr before being recorded for 10 min in each test condition: first in a dish that does not contain prey, to observe *spontaneous* behaviour, and then in the presence of prey to observe *evoked* hunting behaviour (see Section 4.B). Video recordings were then analysed using custom-written tracking software (see Section 4.B). We defined and detected hunt events as blocks of consecutive video frames showing a persistent convergent saccade (≥ 45°) [5, 6, 33, 43]. We will refer to the number of detected hunt events as hunt frequency, as all counts were obtained from recording sessions of equal duration (10 min).

For the analysis of behavioural data we developed statistical models of behaviour and took a Bayesian approach to inferring parameters (see Section 4.F), which utilizes a combination of data and prior insights to obtain updated posterior insights of a model’s parameters. Unlike less informative methods that provide point values of maximum likelihood model parameters, our conclusions are drawn by comparing distributions of these posterior parameters between statistical models derived from the behavioural data of the different rearing groups.

### 2.A Experience increases hunt rate but decreases hunt duration

Rearing conditions may affect the motivation to hunt and/or the ability to detect prey [20, 28]. Although hunting behaviour is always accompanied by eye vergence, eye vergence may also occur spontaneously in the absence of prey. To account for this, we compared evoked and spontaneous eye vergence events (referred to as hunt events from now on) across groups. We find that in all groups the evoked hunt rate is markedly increased compared to the spontaneous hunt rate. ***Figures 2A*** to ***2C*** show that the cumulative distribution function (CDF) of evoked hunt frequency is clearly shifted towards higher rates compared to the respective spontaneous hunt frequency CDF of each group. There were examples of larvae, across groups, whose evoked hunt frequency was less than their spontaneous one ***Figure 2D***. This reduction could simply be due to unobserved hunt events that occurred outside the recording system’s central region of interest, or it may suggest prey-induced inhibition of hunting behaviour in some individuals.

**Figure 2.**
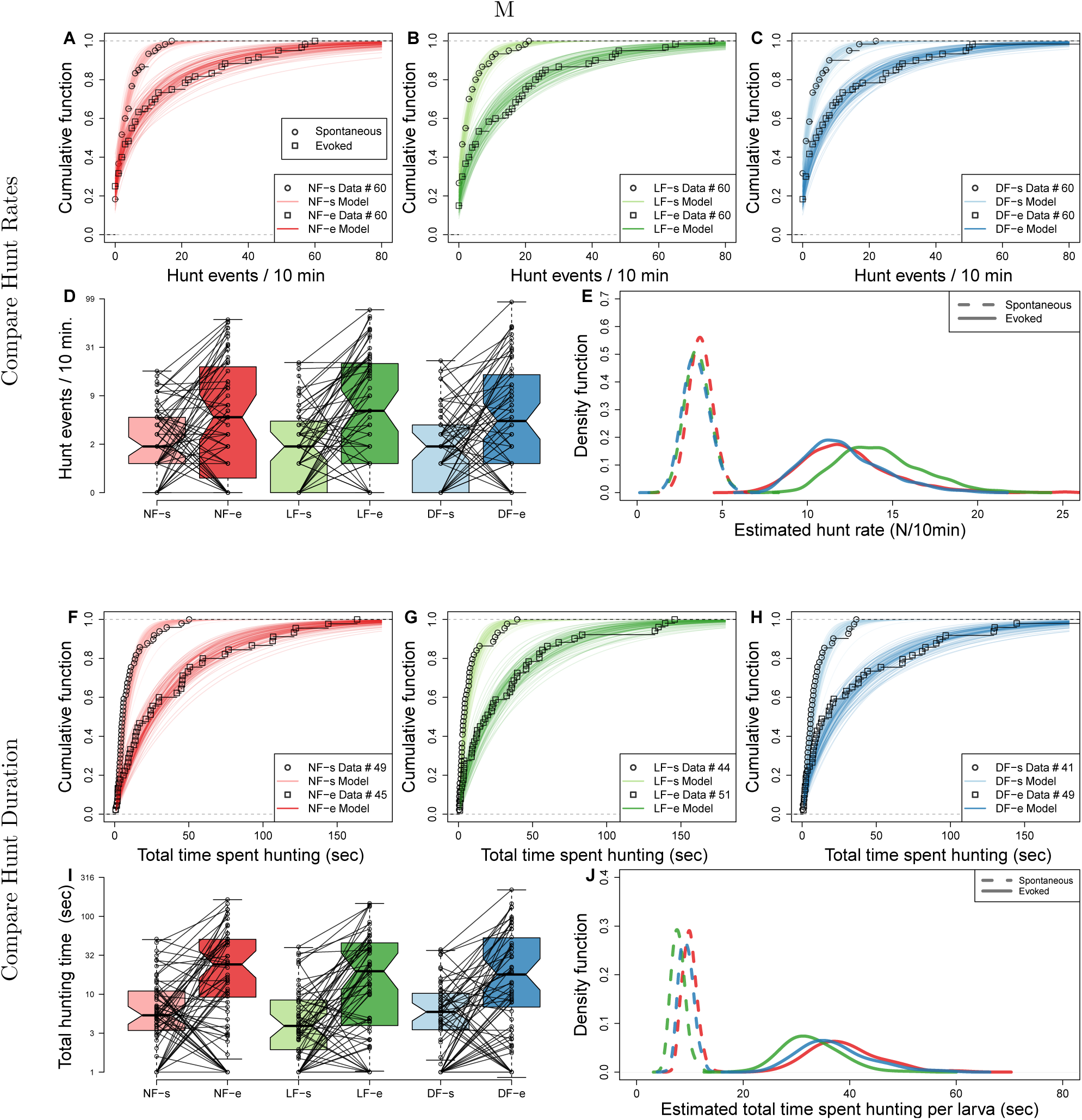
Experience increases hunt rate but decreases total time spent hunting. Hunting effort of each group is characterized in terms of the distribution in number of hunt events and total hunting duration recorded from the 10min behavioural recordings. The two test conditions, in the absence and in the presence of prey, are modelled separately to evaluate *spontaneous* (s) and *evoked* (e) hunt events, respectively. **(A,B,C)** Cumulative density function (CDF) of hunt event counts per larva (open squares) reveals that hunt-event frequency increases across groups once prey is added (NF, not-fed; DR, dry-fed; LF, live-fed). A negative binomial distribution was used to model hunt-frequency data. **(D)** Box plots showing number of hunt events per larvae indicate similar spontaneous and evoked counts in each feeding group. Connecting lines indicate that for most larvae the number of hunt events increases on addition of prey. **(E)** The distribution of estimated mean hunt-rate for each group, as inferred from the models’ parameters, confirms that hunt-rates increase from spontaneous (dotted lines) to evoked (solid lines) conditions. All groups (indicated by line colour) show very similar mean spontaneous hunt-rate, but the LF exhibit a higher mean evoked hunt-rate than the NF and DF groups. Mean estimated group hunt-rate (events/10min) for spontaneous/evoked (LF: 3.6/13.3, NF:3.8/11.9, DF:3.5/12.1). **(F,G,H)** Cumulative function plots showing the total time spent hunting under spontaneous and evoked conditions. Open squares show recorded data and lines indicate 30 cumulative density functions drawn from similar statistical model as in (**A,B,C**,see Section 4.H). All groups show an increase in total time spent hunting when prey is added. **(I)** Box plots showing the amount of time spent hunting increases from spontaneous to evoked test conditions for most larvae. **(J)** The estimated densities of mean hunt-duration of each group, as inferred from the model, clearly show that on average larvae time spent more time hunting in evoked conditions (solid lines) than in spontaneous (dotted lines) conditions. Although DF and NF distributions look identical, the model reveals a noticeable shift towards shorter hunt-event durations for LF larvae in both evoked and spontaneous conditions.

To statistically compare hunt-rates between groups we modelled the data using a negative binomial distribution (see Section 4.G). The resulting model distributions appear to capture our empirical distributions of hunt-frequency very well; the lines in ***Figures 2A*** to ***2C*** indicate 200 CDFs drawn from the negative binomial distributions overlayed with the hunt-frequency data which they model (open squares). The inferred hunt-rate parameter distributions, shown on ***Figure 2E***, reveal that there are no difference in spontaneous hunt-rates between the three groups. The evoked hunt rates for all groups are clearly distinct from the spontaneous rates, and the evoked hunt rates are similar in the DF and NF groups. However, we observed a modest increase in estimated evoked hunt rate distributions for the LF group compared to the two control groups. The mean estimated spontaneous/evoked hunt rates are, LF:3.6/13.3, NF:3.8/11.9, DF:3.5/12.1. These findings suggest that the motivation to hunt may be increased, albeit modestly, by prior experience of live prey. To examine this in more detail we examined hunt durations which may also reflect differences in motivational state.

We measured the total amount of time spent hunting by individual larvae from each group in each test condition and conducted a similar analysis as above, again utilizing the negative binomial. To estimate hunt duration we modelled the total number of video frames per recording that were part of a hunt episode as random events (see Section 4.H). The hunt-duration data show that the total amount of time most larvae spent hunting is noticeably increased in the presence of prey across rearing groups. This is clearly reflected in data and model CDFs of total time per larvae of ***Figures 2F*** to ***2H***. However, when comparing the inferred hunt-duration distributions between groups we find that the LF group is slightly less in both spontaneous and evoked conditions compared to the control groups. Thus, larvae in the LF group exhibit a modest shift to higher distribution of hunt rates but a shift to a lower distribution of total amount spent hunting implying a shift towards shorter individual hunt episodes in the LF group.

To verify this, we examined the duration of all detected hunt events in each condition (699 spontaneous and 2578 evoked), see ***Figure S1***. Although the distribution of episode duration is rather wide in both spontaneous and evoked test conditions across groups (95% of data between 0.5-5 seconds), the mean episode duration is indeed shorter for the LF group than controls, in both spontaneous and evoked conditions.

### 2.B Capture success increases with experience

We next examined whether experience affects capture success. The outcome of 1739 hunt events, in which larvae were clearly involved in following prey, were scored while blind to rearing group (see Section 4.I). As shown on ***Figure 3A***, the success rate for the LF group was the highest at 32%, with 21% for the NF group, and 18% for the DF. We then proceeded to statistically evaluate the likelihood of this outcome by inferring the joint probability density of successful prey capture and hunt-rate (see Section 4.I). For this, we employed a combination of the earlier hunt-rate negative-binomial model and a binomial distribution model in order to infer the probability of success *p* by combining the data on the number of capture successes for a given number of hunt events (Section 4.I). ***Figure 3B*** shows that the estimated probability of a successful hunting event is clearly increased in the LF group compared to the NF and DF groups. Consistent with the ordering seen in ***Figure 3A***, the NF group shows a modest increase in success probability compared to the DF group although there is a region of overlap between the two distributions, indicating that roughly half of the larvae in the DF group have the same success probability as larvae in the NF group.

**Figure 3.**
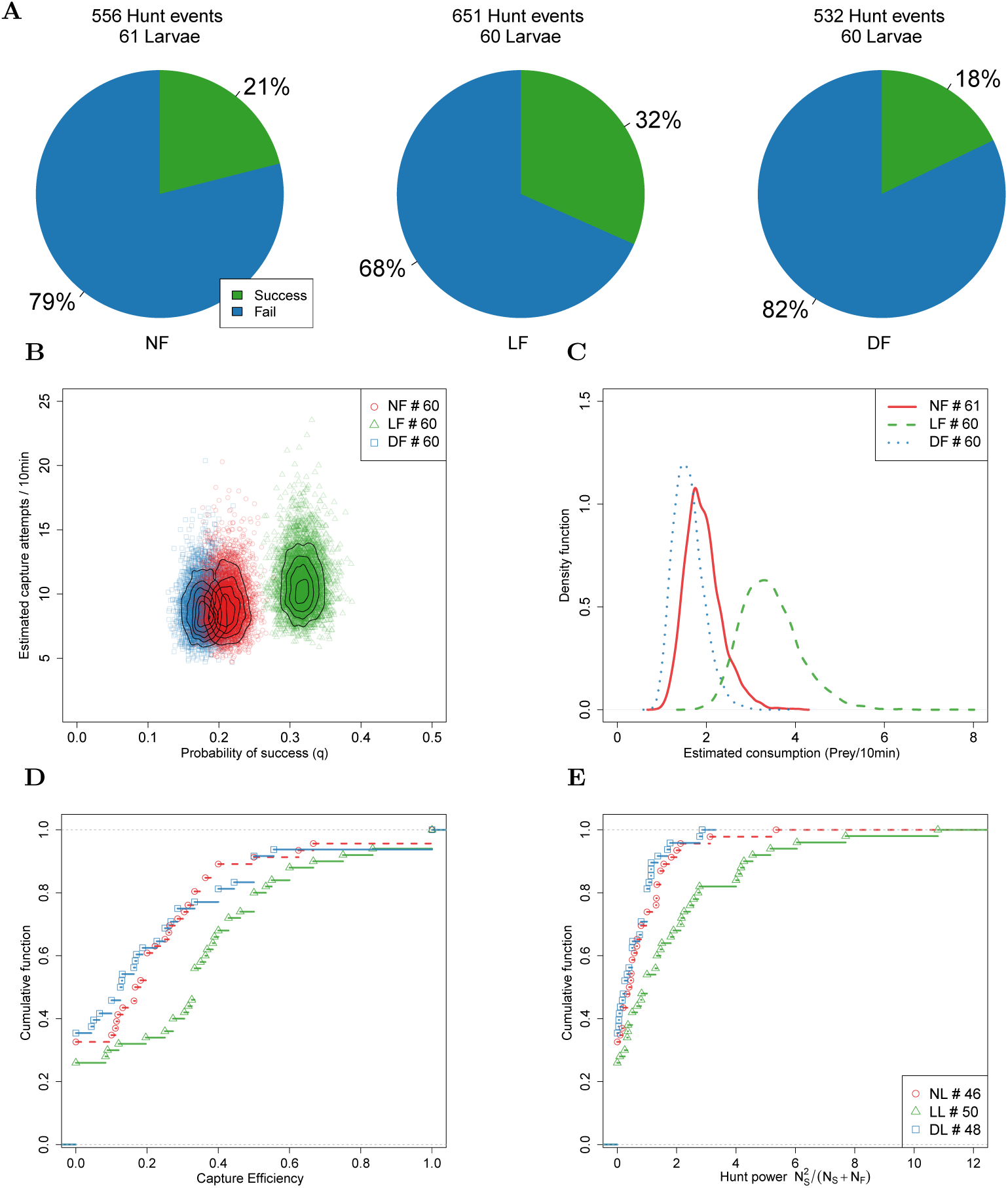
Capture efficiency is experience-dependent. **(A)** Proportion of successful and unsuccessful hunt events in each feeding group based on manually labelled outcomes of hunt episodes. Ranking of proportion of successful hunt events: LF*>*NF*>*DF. **(B)** We model the distribution of mean probability of success against hunt rate (see Section 4.I). Each point represents a likely value for the estimated quantities, and contour lines indicate their distribution. The likely distribution of success probability for the LF group has a distinctively higher mean (¿ ×1.50) than the NF and DF groups. Consistent with ***Figure 2E*** the model predicts a modest increase in hunt rate for the LF group, although the distribution of likely hunt rates for the three groups overlap considerably (estimated mean hunt rates LF: 10.9,NL: 9.2, DL: 8.9). **(C)** Combining estimated hunt and success rates we plot the probability density function (PDF) of likely consumption per group and find that the LF group’s consumption is almost double that of the NF and DF groups. **(D)** CDF of hunt efficiency in terms of fraction of capture successes against capture attempts, shows a general shift rightwards for LF, meaning fewer lower performing larvae as a result of experience. Approx. 40% of LF larvae have efficiency above 0.4, while this is just 0.2 for the DF and NF groups. **(E)** We define a hunt power index (HPI), as the product of efficiency and number of captured prey, in order to account for the number of captured prey in the scoring of hunting ability. A cumulative HPI distribution for each group shows a general shift in the slope, suggesting experience enhances HPI for most larvae, with a subset of 20% of individuals in the LF group having an HPI higher than top performing DF, NF larvae.

We estimated a distribution for the consumption rate per group using the product of hunt-rate and probability of success (***Figure 3C***). Consumption estimates reveal that larvae from the LF group have a mean consumption rate that is almost double that of the NF and DF groups. We find that DF and NF consumption distributions overlap considerably, with mean consumption approximately 1.8 prey/10min, while LF’s consumption distribution is centred at a mean rate of 3.6 prey/10min. Although both DF and LF groups received nutrition during rearing, DF show the lowest consumption performance among the three rearing groups. Our finding that probability of success and consumption rate are increased in the LF indeed suggests prior experience of live prey increases the hunting success rate. Next, we sought to determine the distribution of hunting-ability among the larvae of each group.

We define capture efficiency as the fraction of capture successes over the total number of capture attempts (*N*_Success_*/N*_Total attempts_) for each larva, and plot the empirical CDF in Figure 3D. This distribution suggests that experience has increased consumption rate in the LF group by reducing the number of individuals in the lowest range of hunt efficiency. However, this efficiency measure does not fully reflect hunting performance, because it does not consider the total number of hunt events of each larva, e.g. a larva succeeding in 1 out 2 attempts appears equivalent to a larva that succeeded in 5 out of 10 attempts. For this reason we defined a hunt power index (HPI) to factor each larva’s capture efficiency with the number of captured prey. The empirical CDF of HPI shown in ***Figure 3E*** reveals that (20%) of larvae in the LF group have a hunt power that exceeds the highest HPI for larave in NF and DF groups. Additionally, comparing the slope of the LF group’s CDF to controls we find hunting performance has been modified broadly across the group and not in a discrete subset of individuals.

### 2.C A fast capture swim is the strategy of success and experienced larvae employ it more often

Selecting the appropriate capture strategy for the type of prey, and executing it with accuracy and precision, will strongly affect a predator’s capture efficiency [12]. At the final stage of the hunting sequence zebrafish larvae execute a capture manoeuvre, see ***Figure S15***. These capture manoeuvres vary in apparent vigour and distance to the prey from which they are initiated, and can be divided into at least two types based on tail posture and swim speed classification [31, 34, 43]. We therefore explored whether experience modifies capture strategy and whether this associates with the increased hunting performance of the LF group.

We began by manually classifying capture manoeuvres as either slow or fast, while blind to the rearing group. ***Figures 4A*** to ***4C*** reveal that in all three groups the majority of successful hunting episodes involve fast capture swims and that the majority of failed hunting episodes involve slow capture swims. Although slow capture types form the majority of capture swims in all groups, the LF group, which has highest percentage of capture success, also has the largest proportion of high-speed captures (≈ 41%LF, ≈ 21%DF and ≈ 28%NF), and the highest ratio of fast captures over slow captures in successful episodes (approximately ×5 LF, ×1.69 DF and ×3 NF).

**Figure 4.**
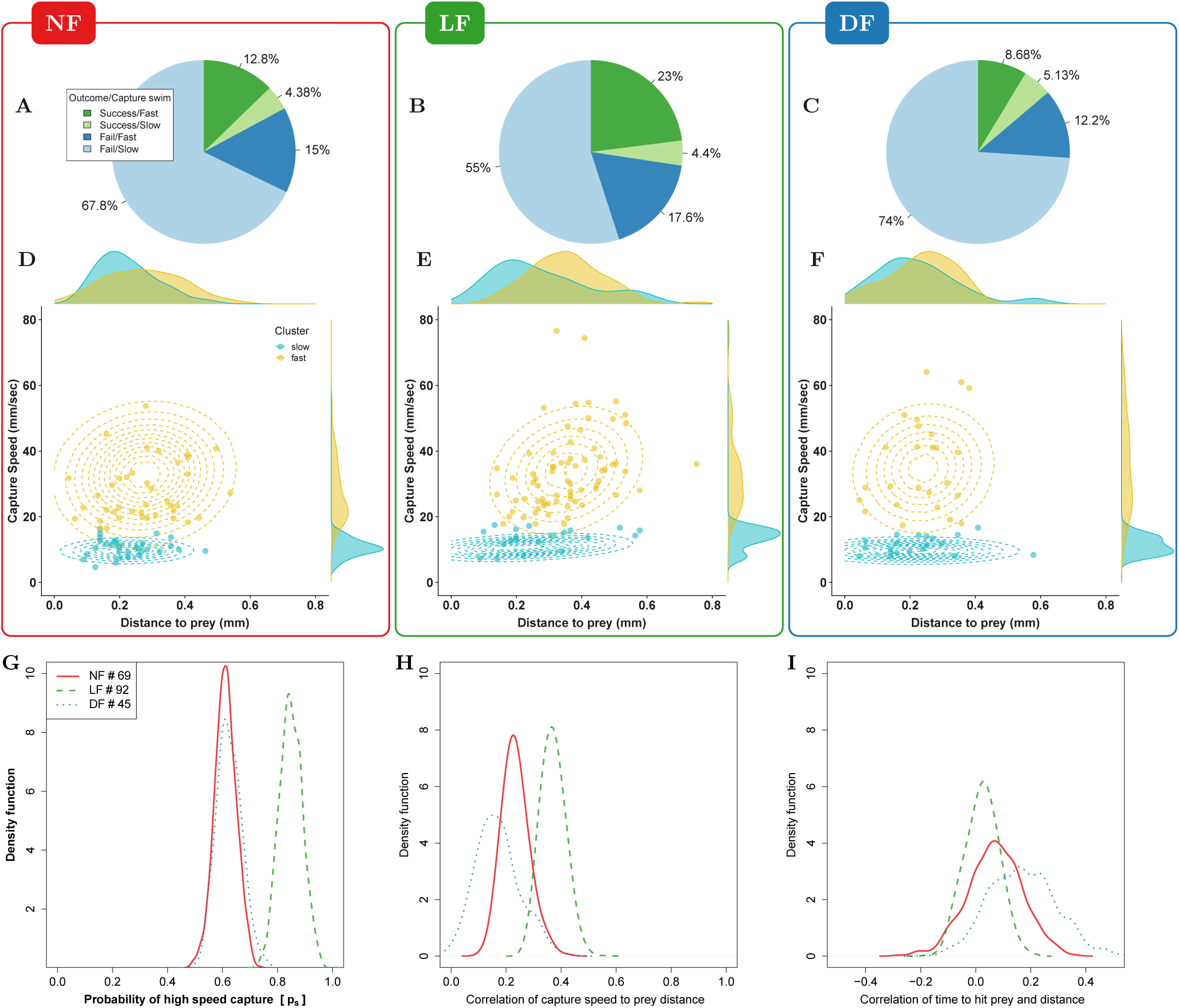
Experience-dependent adaptation of capture swims. **(A,B,C)** Manual labelling of capture types: The majority of successful hunting episodes involve a fast capture swim and these are more frequently observed in LF group. **(D,E,F)** Peak capture-swim speed and prey-distance from successful captures only, clustered in fast/slow capture swim (yellow/cyan) using a mixture of two Gaussians model (shown by contour lines). Shape of fast-capture Gaussian model (yellow contour) suggests that peak capture-speed is modulated by prey-distance in the LF group (see Figure S3). The marginal density plots of prey-distance reveal that slower captures are initiated from closer to prey than fast captures. An increase in the number of successful captures executed from beyond 0.4mm is evident for LF in comparison to DF,NF. **(G)** The fraction of points that are likely to be classified as high-speed capture swims (yellow cluster) by the model is higher in the LF group than in the other groups, and is in agreement with our labelled results (**A-C**). **(H)** Bootstrapped (80% data) distribution of Pearson’s correlation between all capture speed and distance to prey data points shows that all groups increase peak capture speed with distance from which the capture swim is initiated. This relationship appears stronger in the LF group and also agrees with the model’s ellipsoidal shape for fast-cluster (see **E**). **(I)** Density from bootstrapped (80% data) Spearman’s correlation coefficient between the distance to prey and the time it takes to reach the prey in fast-capture swims alone. The time to reach prey does not strongly depend on the initial distance, likely due to the above adjustment of speed with distance, and this is most clear for the LF events. A scatter plot of the capture travel time data is shown on ***Figure S17***.

Collectively, these results indicate that successful captures are more likely to result from fast capture swims, and that fast capture swim are employed more frequently and more effectively by larvae in the LF group. However, the above breakdown was based on a subjective estimation of capture speed from which it is difficult to establish reproducible and accurate classification criteria.

For an objective classification of capture swim types we employed a statistical approach that clustered captures based on their speed and distance from prey (see Section 4.K). Capture speed was measured as the peak speed (mm/sec) during the last motion bout in the hunting sequence prior to a successful prey capture (“capture bout”), and prey distance was measured from the tip of the mouth point prior to capture bout initiation (see Section 4.K). The speed and distance data points were then clustered based on a model composed of a mixture of two joint-normal distributions (see Section 4.K). We limited our analysis to successful hunting routines only (*n*_NF_ = 69, *n*_LF_ = 92, *n*_DF_ = 45), because in these instances the aims and outcomes of the observed behaviour were unambiguously the same. In contrast, comparing between failed hunting sequences would be less straightforward, as failures can occur for many reasons; larvae lose track of the target, abort hunting sequences for no obvious reason, and furthermore, the intended target during failed hunt-sequences is not always obvious.

The clustering results according to capture type are shown on ***Figures 4D*** to ***4F***, coloured according to cluster membership along with the respective contour lines of the joint-normal distribution model for each cluster. In ***Figure 4G*** the probability density of a data point being classified as a fast capture swim confirms that fast capture swims are more likely in the LF group than in the control groups. The data also informs the model’s mean capture-speed and mean prey distance for each cluster, fast or slow. We find that in general the cluster centres of fast capture swims are located at longer distances from prey than the slow cluster centres, while the fast capture swims of the LF group tend to be initiated further from prey than controls.

### 2.D Capture swim speed and distance to prey become more correlated with experience

We next examined whether distance to prey and capture speed are related and whether the relationship is modified by experience. A relationship between distance to prey and the peak capture speed is apparent in the shape of the fast cluster model, at least for the LF group ***Figure 4E***, (captured in the models’ covariances across groups ***Figure S3***). To verify this we measured the Pearson correlation between all prey-distance and capture-speed data points and plotted the distribution of coefficients obtained by repeatedly sampling the correlation (*n* = 10^3^) from a random subset of 80% of data points (bootstrapping). ***Figure 4H*** confirms that a correlation between these variables exists overall and it is stronger in the LF group. Consistently, we find that the covariance of the fast-cluster model also relates capture speed and distance to prey, ***Figure S4***. Overall our findings demonstrate that the frequency of fast capture swims, the distance-to-prey at capture initiation, and the speed-distance correlation increase as a result of rearing with live prey.

For successful capture, the timing of mouth opening needs to be synchronized in relation to prey proximity such that prey enters the mouth cavity either via suction or engulfment [12, 16, 24, 31]. One way of achieving precision in this timing would be to maintain a consistent distance to prey from where to execute capture swims [12]. Alternatively, the capture speed could be adjusted with prey distance in an effort to maintain the timing from capture initiation to reaching prey constant. To evaluate this hypothesis we calculated distributions of Spearman correlation coefficients, by bootstrapping on 80% of the data points, to reveal if there is relationship between the observed time to reach prey and the prey distance while preventing correlations being dominated by the absolute scale of the variables. Figure 4I suggest that time to reach prey does not vary with distance travelled during the capture swim, and this is more clearly demonstrated in the LF group’s fast capture swims. Thus, the ability to adjust capture speed as a function of prey distance is modified by experience. Next we examined whether the accuracy with which larvae re-orient towards prey during pursuit may also contribute to the LF group’s hunting efficiency.

### 2.E An off-axis approach strategy develops through experience

A large turn that re-orients a larvae towards prey commonly occurs at the beginning of the hunting sequence [6, 43, 51] (see ***Figure S15***). This initial turn is proportional to prey azimuth [43, 51] (see ***Figure 5A***). We examined if the accuracy of this response is modified with experience, limiting our analysis to successful hunting for the reasons outlined above.

**Figure 5.**
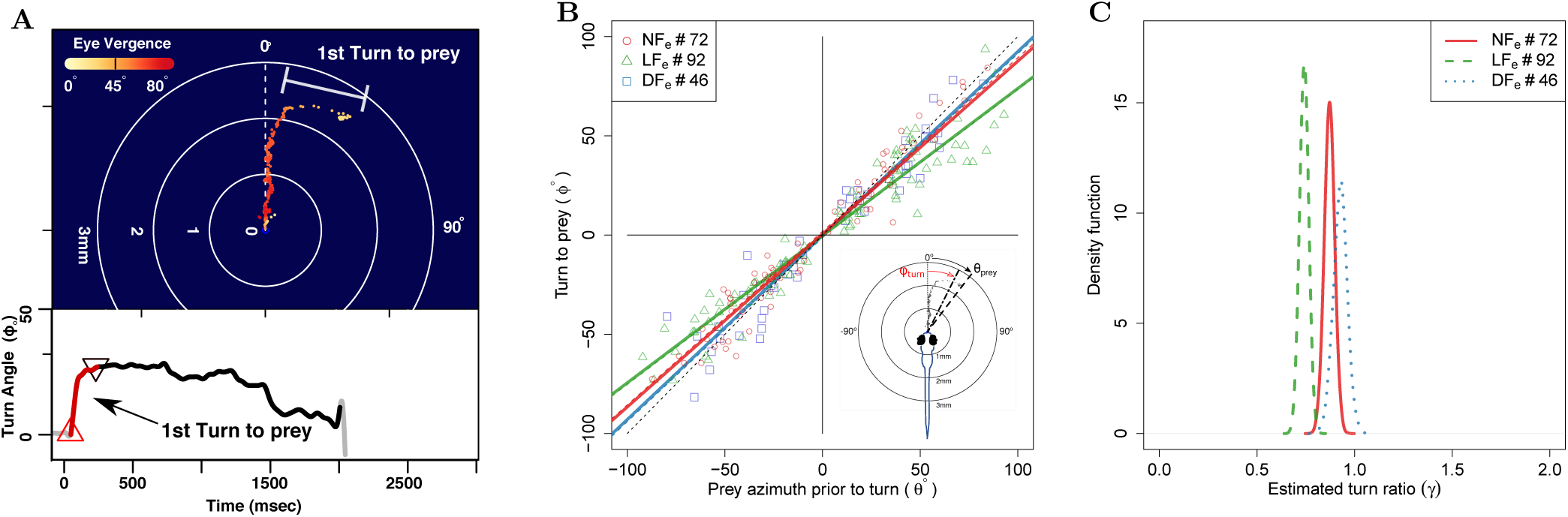
The angle of first turn-to-prey is experience-dependent. **(A)** Hunting sequences begin with eye-vergence and a turn bout that re-orients larvae towards prey. We isolate the initial re-orientation towards prey as shown in the example hunting trajectory, which is colour-coded according to eye vergence. Prey is located in the centre of the figure. A large amplitude turn that re-orients the fish towards prey coincides with an increase in eye vergence angle. Inset shows turning over time, with red highlighting the extracted first-turn behaviour and light grey indicating post-capture turn. **(B)** Prey azimuth vs magnitude of re-orienting first-turn along with linear regression lines for pooled data points separated per group. Hunt events from the LF group show the highest deviation from dotted line, which indicate the slope of turn angles that would precisely align larvae with prey. **(C)** The inferred slope density from the linear statistical regression model reveals that first-turn behaviour in hunt events from the LF group have a lower slope that is distinct from the NF,DF turn events. The ranking in turn-ratios goes LF¿NF¿DF

We first established that prey detection occurred over similar angles across the three groups prior to the initial turn. The distributions of the initial prey-azimuths in successful hunt episodes appear bimodal, see ***Figure S6***, with prey detection from all groups being more likely to occur in response to prey located 35°-50°on either side of the midsaggital axis. A scatter plot of turn response to the initial prey azimuth suggests that these two quantities are proportional ***Figure 5B***. The ratio of these two quantities (“turn-ratio”) is unity when larvae accurately re-orient towards prey. We found that the data pooled from recorded hunt episodes across larvae gave a mean turn-ratio (*µ*± SEM) 0.80 ± 0.029 for LF group, and this was different to DF 0.98 ± 0.05, and NF 0.93 ± 0.03 (see ***Figure S5*** for histograms). To evaluate potential differences between group data statistically we used a standard linear to model turn-ratios by the inferred slope (see Section 4.J).

The distributions of likely turn-ratios estimated by the model shown on ***Figure 5C*** confirm that the initial turns recorded in the LF group are distinct in undershooting prey azimuth; the slope of the regression model gives a mean 0.73 for LF, 0.87 NF, and 0.92 for DF. This undershooting behaviour recorded from the LF group is consistent with previous reports on the first-turn-to-prey of larvae that had also been reared with live prey (Paramecia) [8, 43, 51]. Paradoxically, hunt events from the NF and DF groups display initial turns that do not consistently undershoot, but rather align larvae closer to the prey’s azimuth Figure 5C. We conclude that undershooting on the first turn to prey is a kinematic adaptation of the hunting sequence that result from rearing with live prey. We next examined whether experience is required for linking the initial turn behaviour with fast capture swims, which were more frequent in the LF group.

### 2.F Experience combines off-axis approach and fast capture swim strategies

The LF group exhibits a higher frequency of fast capture swims and a stronger tendency to undershoot in their initial turn to prey than controls, but whether these two behaviours are combined within individual hunt events remains unknown. Fast capture swims and undershooting could be used independently of one another or they could be linked either via a learned or innate association. In the case of an innate relationship, which is immutable by experience, undershooting would lead to fast-capture swims regardless of rearing group. Alternatively, if the association between these two behaviours can me modified by experience, the combination of undershoot and fast-capture swim should be unique to hunt events of the LF group.

The relationships between turn-ratio and capture-speed for each successful hunt episode are shown on ***Figures 6A*** to ***6C***, along with densities for each of the two clusters (fast/slow) on the plot margins. An association between fast capture swims and turn ratio would manifest as a leftward or rightward shift in the densities shown along each plot’s top margin. A leftward shift, indicating undershooting (turn-ratio ¡ 1), is visible in the LF group’s turn-ratio density of the fast capture swims (yellow) on ***Figure 6B*** while no obvious relationship between turn-ratio and capture speed is seen in the fast capture swims of NF and DF groups ***Figures 6A*** and ***6C***. Therefore, the fast-cluster data-points from successful capture episodes show a clear bias towards one side of the turn-ratio only in the LF group.

**Figure 6.**
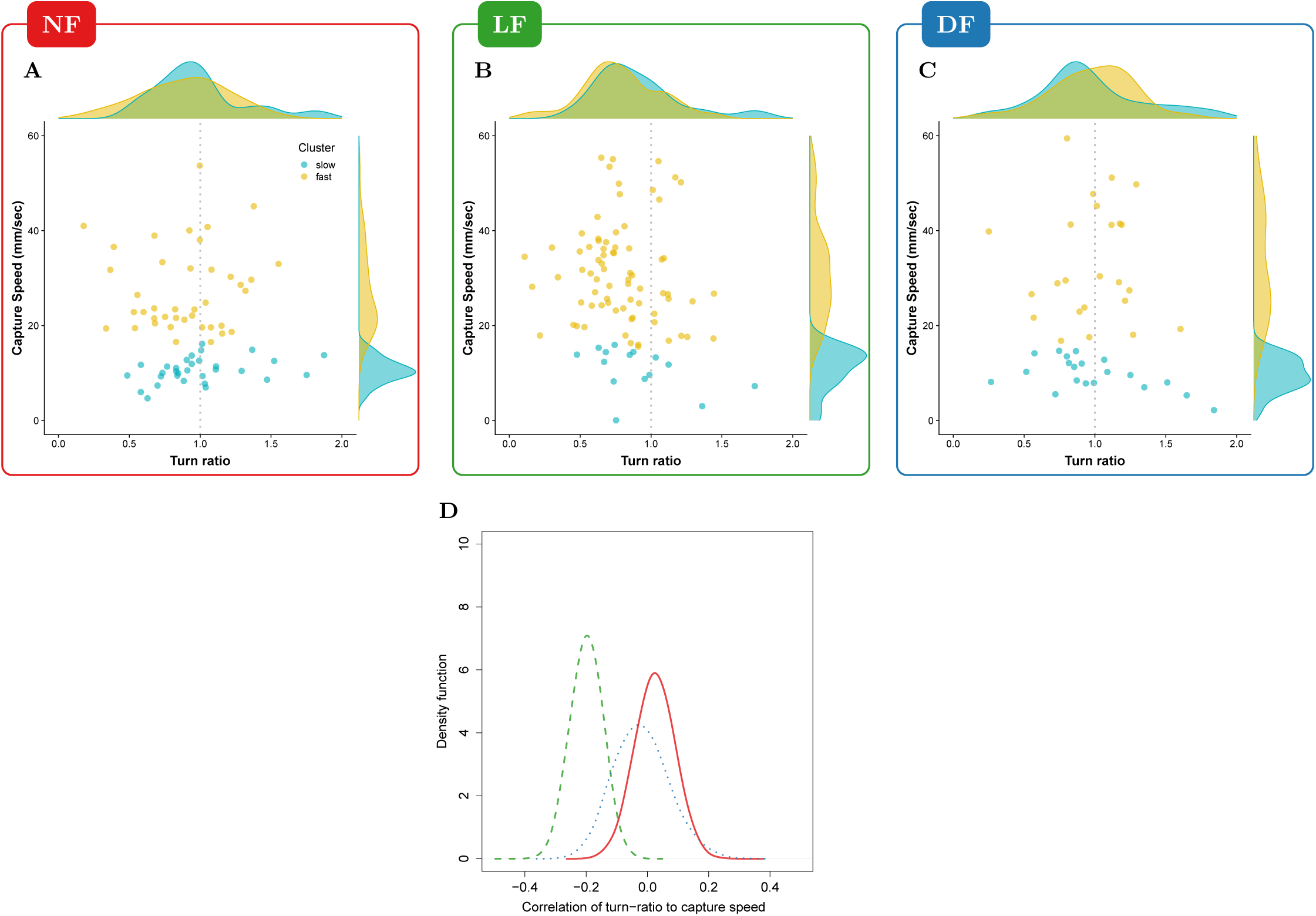
Experience associates undershooting to fast capture swims. **(A,B,C)** Pooled data points across larvae on capture-speed and turn-ratio. The densities shown along the top margins suggest that fast captures (yellow, see Figure 4) are biased towards low turn-ratios (undershooting) in the LF group, while no such preference is seen in the NF and DF groups. **(D)** Distribution of bootstrapped (80%) Spearman’s correlation coefficient between turn-ratio and capture-speed; Hunt events from LF larvae show a relationship between low turn-ratio (undershoot) on their first-turn to prey with the speed of the final capture swim. This relationship is not innate but rather driven by experience because although DF and NF larvae may occasionally undershoot on their first-turn to prey, this turn is not associated with a fast-capture swim.

To evaluate the evidence for a correlation between turn-ratio and capture speed we calculated densities of their correlation by taking multiple random subsamples (80%) from all successful hunting data for each group and calculating Pearson’s correlation coefficients for each subsample. ***Figure 6D*** shows correlation densities clearly associating undershooting and capture speed in the hunt events of the LF group while no such clear relationship is evident for the hunt events of the control groups. Thus, undershooting on the first-turn-to-prey and fast capture swims are correlated in the LF group but not in inexperienced larvae. This suggests that experience is not only required for modifying these two components of the hunt sequence but also for chaining the modified components together into a sequence.

### 2.G Typical hunting behaviour is distinct in experienced larvae

Our analysis so far has found that the experienced group’s hunting routines are different to control groups’ in terms of turn-to-prey and final capture swim. However, these observations may not be representative of typical larval behaviour within each group, since the majority of successful hunt-events analysed could be originating from an atypical minority of larvae. In this section, we go beyond comparing pooled hunt-episodes and characterize the prevalent hunting behaviour of each group’s larvae.

We summarize typical hunting behaviour of each group in a hierarchical statistical model that builds a model of group behaviour by aggregating estimates of mean hunting behaviour from each of the group’s larvae (Section 4.L). Consistent with our earlier analysis, which utilized successful hunting episodes alone, we draw our estimates for the mean hunting behaviour of each larva based on its turn-ratio, capture-speed and distance to prey. We then compare hunting behaviour between groups in terms of distributions in the space defined by the model parameters that reflect each group’s hunting behaviour. ***Figure 7A*** shows a 3D rendering of the resulting posterior parameter distributions per group, according to which the LF group (green) has higher overall capture speed and distance, and lower turn ratio (see ***Figure S8*** for a more detailed view of parameter space). This result is consistent with our earlier analysis of pooled hunt events (***Figures 4*** and ***5***). Nevertheless, with this group model we can confirm that experience has indeed affected the hunting behaviour of the larval population and we can also obtain the estimated mean hunting behaviour of each larva.

**Figure 7.**
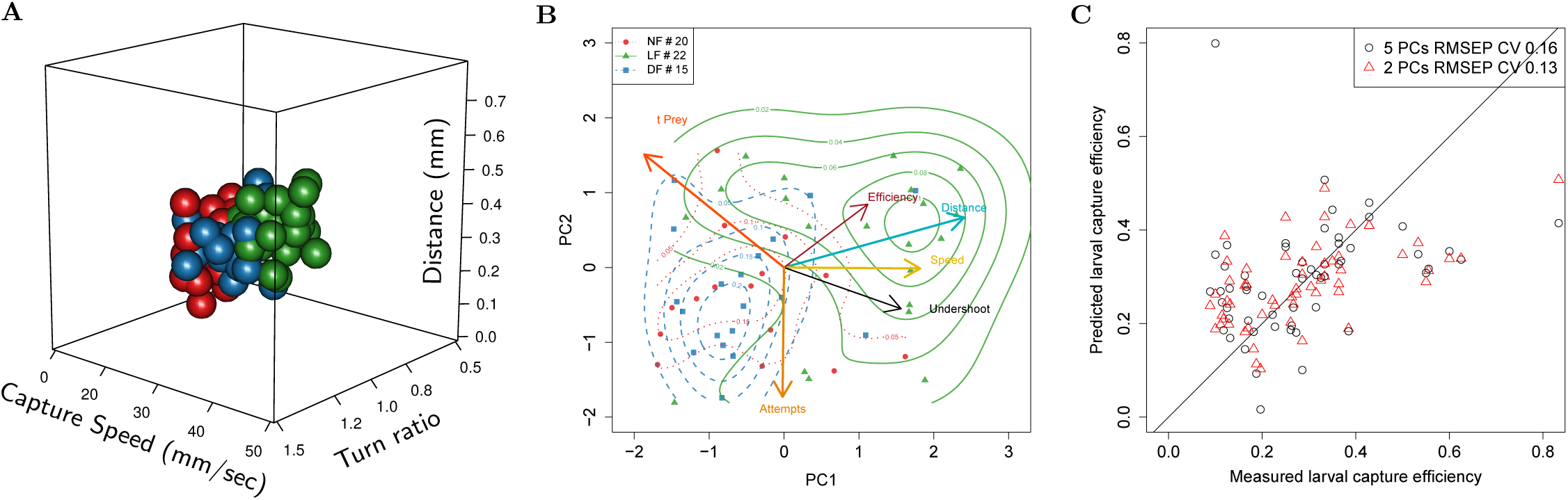
Typical behaviour of experienced larvae is distinct and relates to their increased capture efficiency. **(A)** Estimated distribution of mean hunting behaviour based on capture speed, distance to prey and initial turn for each group according to hierarchical model of group behaviour. The LF group’s model shows that successful hunt episodes are more likely to have capture strikes initiated from further away to the prey than control groups. This is also true for turn-ratio, overall reflecting an expectation that larvae in the LF group are more likely to express undershoot and higher peak capture speeds. **(B)** PCA in larval hunting behavioural space showing the axes for hunting behaviour and the position of individual larva across groups as seen from the first 2 PCs. Larva behaviour was estimated by the model and associated with capture efficiency (min. 5 capture attempts) in this PCA space. Distributions per group (contour lines) confirm separation of the LF group from controls and shift toward region pointed to by the efficiency axis (red). Members of the LF group are shown to be more efficient hunters, they undershoot on their first turn to prey (black), and execute fast capture swims (yellow) from a relatively longer distance to prey (cyan), thus suggesting a relationship of these behaviours with hunting efficiency. **(C)** Linear regression model shows that 2 principal components of a larva’s mean behaviour, as in **B**, can be used to predict its capture efficiency.

### 2.H Capture efficiency is predicted by the combination of off-axis approach and fast capture swims

Next, we provide evidence that the adaptations observed in the hunting routines of the experience group can partly explain their increase in capture success. For this analysis we only included larvae for which we had at least 5 capture attempts in order to reduce the error in estimating the mean capture efficiency per larva. We then compiled a vector for each individual that contained its estimated capture efficiency along with the number of capture attempts and the three values (capture-speed, distance to prey, turn-ratio) that characterize its hunt-behaviour, as derived by the earlier model. The relationship of these variables was examined by doing principal component analysis (PCA) on a matrix composed of the above larval behaviour vectors across groups. In ***Figure 7B*** we show the multidimensional space defined by six hunting variables as viewed from the reduced space of the first and second principle components (PC). These two PCs capture approximately 62% of the total variance and approximately 31% of variance in capture efficiency. Consistent with our group model in ***Figure 7A***, we find that LF larvae occupy a distinct position in this space that separates them from the control group NF/DF larvae. The LF region is pointed to by axes of capture speed, prey distance and undershoot, as well as increasing efficiency; larvae from control groups occupy the opposite region in this behavioural space which is in the direction of lower efficiency.

This coarse alignment of efficiency with the axes for capture-speed, distance and undershoot indicates that hunting efficiency may be related to, and possibly predicted by these specific hunting behaviours. To examine this further we employed principal components regression [54] and assessed the prediction capability of linear models that could utilize any combination of PCs as factors. The maximum cumulative percentage of efficiency variance that can be explained with two PCs, given the factors we used on ***Figure 7B***, was found to be ≈ 31%, and the coefficient of variation (CV) in the root mean squared prediction error (RMSEP) was *CV* = 0.1335. In comparison, using all five PCs increased the efficiency variance explained to ≈ 36%, but also the RMSEP CV= 0.1633. The ability of these PC efficiency prediction models are shown on ***Figure 7C***.

We have therefore obtained evidence that capture success relates to, and can be partly predicted by utilizing covariates of hunting behavioural measurements within the first 2 PCs. Additionally we found that these 2 PC are loaded with combined behaviours (as in ***Figure 7B***), revealing that increasing efficiency relies on adapting behaviours in a coordinated manner, as is the case for example between capture-speed and prey distance.

## 3 Discussion

We have shown that first-feeding zebrafish can improve their hunting performance by engaging in their natural hunting behaviour. This can be observed as early as 7 dpf, and to our knowledge, it is the first time experience has been shown to affect hunting ability at that early developmental stage. Our experimental design allowed us to examine the comparative effects of different rearing diets on hunting behaviour, and disentangle the effects of growth and nutrition from those of experience. By introducing novel rigorous methods that accurately characterize and compare the effects that rearing food has on the hunt-effort and the capture success of each group, we were are able to provide the first comprehensive evidence that differences in consumption are mostly due to experience-depended changes in hunting skill, with the least efficient hunters in a population benefiting the most (see ***Figure 3D***). We then focused on identifying the behavioural aspects that lead to increased capture efficiency by conducting detailed kinematic analysis of hunting sequences. Utilizing statistical modelling of behaviour at the individual larva, and the population level, we were able to build a prediction model of hunting efficiency that associates capture success to coordinated adaptations of the hunting sequence.

### 3.A Experience-dependent changes in hunt rate, duration and efficiency

Because all hunting sequences in larval zebrafish begin with eye vergence [5, 6, 43], we could use the rate and duration of eye vergence events to estimate hunt motivation and the ability to perceive and respond to prey. We found that the rates and duration of spontaneous and evoked hunt events were very similar in the DF and NF groups. However, compared to the two control groups there was a modest increase of hunt rates in the LF group only in the evoked conditions. This finding is consistent with previous observations that zebrafish larvae raised with Paramecia performed prey capture behaviours more frequently than those that had not [43], and suggest that prior experience may condition prey response, which would be in agreement with findings in other fish species [see 56].

Indeed, a companion study that uses functional imaging in the zebrafish brain demonstrates that prior experience of live prey leads to a lower threshold for activation of visual areas that trigger capture initiation (Oldfield et al, submitted to Elife, 2019). However, our finding that hunt rates are not elevated in the NF group is seemingly at odds with studies showing that starved larvae are more likely to shift behavioural decisions from avoidance to approach [20] and increase their food intake compared to fed larvae [28]. This apparent discrepancy can be reconciled by the fact that we allow all feeding groups to acclimatize with live prey for 40min-2hr prior to recording in the evoked conditions. This time period has been shown to be sufficient to overcome motivational differences between starved and fed larva, as within 40mins differences in the bout patterns expressed between these groups become undetectable [27].

Despite the increase in their hunt rate, larvae in the LF group tend to spend overall less total time hunting than larvae in the control groups. This decrease is noticeable in both, spontaneous and evoked conditions, and thus it may not depend on the presence of prey. Although, it is still likely that experienced larvae become faster at catching prey and therefore more time-efficient, the evidence here suggests a change in an internal timing process which is responsible for sustaining the state of hunt activity. Indeed, the distributions for the duration of individual hunt episodes are surprisingly similar between the evoked and spontaneous events of each group (***Figure S1***), suggesting that an intrinsic timer is also involved in controlling the duration of each hunt episode. This hypothesis of internal timing of foraging state has only very recently been confirmed [32], and our evidence here suggests that these internal dynamics can be altered by experience.

We then extended our statistical model of hunt rates to include the probability of success of each of the evoked hunt events. This more comprehensive model revealed that the estimated mean prey-consumption rate for the experienced group is approximately double that of control groups. This increase in consumption was largely due to improved capture success rather than increased hunt-rates, because the probability of a successful capture was significantly elevated in larvae from the LF group by ≈ 50% compared to NF larvae, while the difference in the number of capture attempts between the LF and NF groups was more modest (≈ 18%) (see ***Figure 3B***). Thus, the effects of experience on hunting behaviour are mostly seen as an increase in capture efficiency at this stage of development (7dpf). The superior capture efficiency of LF against the NF group is not explained by differences in growth, because larvae in the nutrition control group (DF), which tend to be larger than larvae in the NF group (see S13), have the lowest capture efficiency of the three groups. Therefore the most likely explanation is that superior capture efficiency is caused by changes to the hunting routine.

### 3.B Experience-dependent modification of the first-turn-to prey and final capture swims contribute to capture efficiency

There may be multiple paths to success, but some can be more efficient than others. Accordingly, even when comparing hunting episodes that end up in successful captures, there may be differences in hunting behaviour between efficient and non-efficient hunters. Indeed, by comparing behaviour during successful hunting episodes alone, we were able to unambiguously identify differences in components of the hunting sequence between experienced and control larvae.

We show that hunting sequences of experienced larvae are distinct in at least two respects. Firstly, on detection of prey, larvae from all groups make an initial turn that accounts for a large part of the reorienting response. However, experienced larvae make a first turn that tends to undershoot prey azimuth, whereas larvae in the two control groups make an initial turn that brings them closer to aligning with prey azimuth. Secondly, we find that experienced larvae are more likely to employ high speed capture swims and that these are initiated at greater distances from prey than the control groups. Capture speed and distance are correlated, yet this correlation is highest in the hunt events of the LF group, implying that experience is necessary for tuning the vigour of the capture swim as a function of prey distance. To verify that these behaviours are characteristic of experienced larvae, we embedded larvae of all groups in a principal component space of hunting behaviour and found that larvae from the LF group are segregated against the NF and DF larvae, while NF and DF strongly overlapped.

Using only just two principal components we able to build a linear model that predicts larval hunting efficiency, in terms of the fraction of capture successes over total capture attempts, and therefore show that the identified hunting behaviours are related to the probability of capture success. Neither of the two PCs aligned with a particular hunt behaviour in isolation, but rather each component fused a mixture of first-turn-to-prey, capture speed, capture distance and number of capture attempts. However, why does modification of capture speed or an undershoot turn-ratio increase hunting efficiency?

### 3.C Fast capture speeds from a distance prevent prey escape

Capture probability can vary with prey type as different prey are equipped with different motion and escape patterns [12, 15, 36]. For some evasive prey species utilizing a fast capture speed may be important [16]. There is already evidence that Atlantic salmon, carp (*Cyprinus earpio*) and pike (*Esox IUC US*) larvae improve their capture success by using fast capture-speeds [12, 16]. Additionally, attacking prey from a distance can minimize the chance of being detected by prey and evoking escape responses. Thus, it would be beneficial to adapt the capture strategy based on patterns of prey motion, and specifically develop fast capture swims from distance for particularly evasive prey.

Nevertheless, this strategy poses new challenges because as a larva lunges forward during a capture swim the gape and suction action need to be coordinated such that they occur right in front of the prey [12, 31]. Otherwise, failures occur because the larvae hits and pushes the prey away [12]. Indeed, zebrafish larvae show a fixed gape-cycle [24] and by adjusting swim speed as a function of prey distance, experienced larvae can set the timing of hitting prey according to their gape-cycle. By examining the capture distance-speed relationship in terms of time to hit prey, we found that this was more accurately adjusted in experienced larvae (see ***Figure 4I***). An increase in timing accuracy can facilitate prey capture by allowing the timing of the mouth opening to occur in proximity to prey as the larva lunges forward. However, how can larva perceive their distance to prey, and know they are within the capture distance ?

### 3.D Turn undershoot as an active sensing strategy

We focused our analysis of turn behaviour only on the initial turn towards prey because it amounts to the largest re-orientation following prey detection and so it can be representative of the ability to aim towards prey while being less prone to measurement error. Subsequent turn behaviour follows the same pattern of undershooting prey azimuth, and thus the first turn behaviour we observe is typical of a larva’s re-orienting behaviour overall [8]. We hypothesize that this undershooting of the turns plays a role in enhancing capture success by improving the perception of prey distance. Object size is an important feature for triggering hunting in zebrafish larvae [6]. However, vision in zebrafish is monocular with a fixed focus, and so retinal image size is an ambiguous cue for the true size of an object. A potential a way to resolve this ambiguity is to utilize motion parallax, a monocular depth cue in which we view objects that are closer to us as moving faster than objects that are further away. Through undershooting turns, prey is positioned at some relative angle to the larva’s heading. During larva bouts of forward motion, parallax can provide a measure of prey distance that can be readout through the speed that the prey image is translated across the retina. Such active sensing techniques are employed by insects that have been reported to move in particular angles to objects in order to generate optic flow and discern their proximity [18]. Generally off-axis approach strategies can extend beyond vision and may represent navigational trade-offs. For example an off-axis strategy has been shown to improve object localization in echolocating Egyptian fruit bats, and in these bats, it appears this strategy represents a strategical trade-off between positional accuracy and detection range [59]. In our case, the undershooting strategy may present a trade-off between minimizing the number of bouts required to reach prey and the perceptual accuracy of prey distance.

A prediction arising from the idea that image motion is integrated with retinal image size is supported by empirical evidence on hunting initiation that required the use of specific combinations of stimulus size and speed [5]. We obtained further evidence that prey located off-axis is more likely to be captured, by observing that the distribution of initial prey azimuths across successful hunt episodes (see ***Figure S6***) is bimodal. Although larvae are able to respond to prey in front of them equally well [6, 8], we find that the distribution of prey azimuth is bimodal in successful hunt-events across all groups, and the maxima are located between 35°-50° on either side of the larval heading. Therefore, prey located off-axis at particular angles are more likely to be captured.

### 3.E Learning mechanisms that link behavioural components into a sequence

Experience not only modifies the first turn-to-prey and capture swim, it is also required for linking the modified manoeuvres together. This conclusion is based on the observation that undershooting and fast capture swims are most strongly correlated in larvae raised with live prey, i.e. larvae in the NF and DF groups occasionally undershoot, but this does not correlate well with fast capture swims. This implies that experience does not simply increase the frequency of undershooting and that more frequent fast capture swims are an inevitable consequence. This raises interesting questions concerning the mechanisms of learning that associates these modified manoeuvres. Fast capture swims are more likely to lead to an immediate reward, consumption of food. Thus, fast capture swims may be positively reinforced through operant learning. Undershooting on the other hand does not lead to an immediate reward, so how does experience increase the frequency of this behaviour?

Conditioned, or secondary reinforcement is a learning process in which neutral stimuli (in our case, perception of prey arising from undershooting and motion parallax) that predict primary reinforcers (consumption of prey through fast capture swims) can themselves become reinforcers [30, 55]. However, previous attempts to establish classical or operant learning in larval zebrafish, by pairing visual cues to electric shock, failed to show learning at this early stage [52]. A different, classical conditioning study showed enhanced tail response to the conditioned stimulus after pairing a tactile stimulus on the side of the body with a visual cue of a moving spot [2]. Thus, although in theory a form of arbitrary behavioural chaining could be used to generate efficient behaviour in a wide variety of experienced conditions [19], it would appear here that learning is constrained to expecting specific information about the environment to instruct the parameters of particular behaviours of the developing animal [4]. In this case, early larval hunting behaviour allows them to learn and adapt to the prey available in their environment.

Finally, in our analysis of the distribution of hunt efficiency we found that experience had not made all larvae equally successful hunters, but it was mostly the lowest efficiency hunters that were diminished in the population when compared to controls. Thus, it is very likely that larvae learn at different speeds due to natural variability, but more importantly it appears that a brief, over two days, hunting experience can rescue initially ineffective hunting behaviour.

### 3.F Conclusions

In summary, we have demonstrated that prior experience of live prey modifies and associates components of the larval zebrafish hunting sequence in manner that improves their capture success. Our findings suggest that the ontogeny of hunting behaviour relies on learning by experience to fully develop. Combined with prior attempts that failed to show conditioning in larval zebrafish [52], it appears that learning may be constrained to particular tasks at this early stage, but nevertheless it is sophisticated enough to alter a multidimensional behavioural space so that innate goal-oriented behaviour is improved. Such interactions between an innate behaviour and learning are very common to the ontogeny of behaviour [26] and it is likely that the general learning principles, which allow experience to shape brain development, are conserved across species [53]. This study will pave the way for using zebrafish to study the neural circuits and mechanisms of learning which transform ethologically relevant experience into efficient natural behaviour.

## 4 Materials and methods

### 4.A Rearing

Fertilized embryos under natural spawning were collected at 10.30am from a mass embryo production system (MEPS), where they are developmental stage synchronized within 15 minute collection intervals. These were visually inspected and 17 healthy embryos are selected and placed in each of three 9cm Petri dishes, which were filled with 35ml Daneau and labelled at random to define the rearing group (NF,DF,LF). The embryo dishes were maintained in an incubator under the same conditions. of 28.5 °C in system water (pH 7.3, conductivity 550 *µ*S) on a 14:10 h light:dark cycle. On 1 dpf non-developing embryos were cleared (usually 0-3 dead ones), and from that day on the dish where cleared of debris and had half the water replaced on a daily basis.

Feeding initiated just prior to 5 dpf, around the time when larvae begin to initiate hunting, and continued until the beginning of 7 dpf with a single feed each day between the hours of 3-4pm. Live-fed (LF) received ≈ 200 live rotifers (*Brachionus plicatilis*) (sized between 0.3mm and 0.05), usually in 1-3ml of 2 ppm water, the volume depended on the density of the culture on the day. The Non-Fed group (NF) received 2ppt salt water, to control for treatment and salinity, in a volume matched to one supplied to the rotifer fed dish (LF, 1-3ml) on that day. The health or survival of the NF group was not impacted up to 7 dpf during which time their yolk-sack energy store appears sufficient to keep them healthy, and this is in agreement with studies showing that survival rates remain unaffected even if feeding commences on 8 dpf [25]. The Dry-fed (DF) group receives grounded growth food suspended in the same amount of 2ppm salt water as the one delivered to other groups. The DF food is grounded with mortar and pestle (Sera Micron or Ketting Gemma 75), then suspended in 2ppm water and centrifuged in 800rpm for 20 sec. The suspension, which mostly contains particles smaller than rotifer typical size, is used for feeding such that the visual experience of moving dots is minimized for the DF group but nevertheless they receive a nutrition.

### 4.B Behavioural recording

At 7dpf individual larvae from each group were transferred to 35mm petridishes (5ml Daneau) and were allowed to acclimatize for 40min-2 hours in the test conditions prior to each video recording. The recording protocol’s timeline is shown on ***Figure 1***. Each larvae was recorded in two test conditions. First settled and recorded in a prey-free (empty) arena and then, following the addition of ≈ 30 live Rotifers, larvae are left to settle prior to being recorded with live prey conditions. For the empty test conditions, individual larvae where randomly picked from the 9cm group rearing dish, washed by transferring to a clear Daneau bath, and them they were transferred to their individual 35mm dish containing 5ml of filtered Daneau water, such that to remove any floating impurities that could trigger hunt events. For the live-prey test conditions we added ≈ 30 live rotifers to the same dish as above, and then topped up prey numbers prior to placing the dish on the recording rig. Once the dish was transferred onto the recoding rig, we let it settle for 5-10 minutes before starting the recording software.

Our recording setup is dark-field illuminated via a custom made light-ring composed of 7 infrared (835nm) emitting diodes (VSMY98545) that helped provide high-contrast images of both small prey particles and larval features. These are arranged appropriately such that illumination is uniformly distributed by the converging IR light beams onto the area of the circular 35mm petridish, therefore eliminating light fluctuations and cancelling any directional preferences arising from NIR light sources [21]. We provided a total of ≈ 250mW to the light-ring and tried to keep power low to avoid any thermal currents in the water. Below the arena sits a Chameleon 3 FLIR camera, with 50mm/F2.8 lens kit MVL50M23, supplemented by a 5mm lens spacer (CML05) to provide x3.5 magnification, and an long-pass filter so as to record in IR only. Above the arena sits a frosted-glass on which visible light is diffusely reflected from a directed lab-bench light source, set-up on either side of the rig (see Figure S14). This provided sufficient lighting for the larvae to be able to see and track prey.

Video recording was controlled via custom recording software that allowed us to limit the total recording time to 10 minutes and to minimize video data, by not recording when the larva is not within a central region of interest (ROI). This was set to a 25mm diameter circle in the centre of the 35mm circular petri-dish. Behaviour was recorded at 410 images per second with a resolution 640×512. Raw image sequences were then converted into compressed video files using the highly efficiency H.264 compression codec. Recording events were automatically triggered when an object of sufficient size (¿ 120px area) entered the central ROI, thus ignoring behaviour near the edges of the arena. Each triggered recording event was set to have a minimum duration of 30 sec., and an initial recording event is automatically triggered at the start of each experiment, even if the larva is not within the ROI for this event. This initial short clip allows us to automatically estimate and verify the number of prey at the start of each experiment. If a larva was still within the ROI at expiration of the 10 timeout period, the recording time was automatically extended up to a maximum of 2min to wait for the larva to exit the ROI. This aimed to avoid the 10min timeout interrupting an ongoing hunting episode. If a larvae triggered no recording events in the empty conditions it was then rejected and replaced by a new one on which the recording protocol was restarted, beginning with settling in empty conditions.

Recording began at 11am and continued throughout the day usually ending around 10pm. During the day, room temperature varied from 21-26 degrees Celsius. Recordings from all groups were being balanced for time of day, to control for circadian and temperature effects, as well as batch variability, by recording the same number of larvae per group on each recording day. Overall our dataset includes 15 batches, in total 180 larva (60 for each rearing group), which were obtained between 16 Nov 2017 and 21 August 2018. Behavioural tracking was conducted offline and was extracted from collected videos using custom video analysis software.

### 4.C Behavioural tracking

#### 4.C.1 Larva body tracking

A background model is computed by employing the OpenCV library’s [10] mixture of Gaussians (MOG) background model on the initial 100 frames of video with a learning rate of 1/400, which is then set to a nominal rate of 1/1000. On each video frame, after extracting foreground objects, we filtered for blob area to identify a rectangular frame region that contains the larva. Because larva heads are very similar, we found that the orientation and position can then be easily and quickly located within this subregion using template matching. We compiled a small (20) library of larva head samples sized 22×33px and replicated each sample across 360 rotations with a resolution of 1°. We then utilized these in our tracking software to do template matching and identify the position and rotation of the larva’s head. The identified template rectangle framed the eyes and swim bladder, and its centre, which was located near the anterior end of the swim bladder, was used as the reference point for tracking the larva’s position.

#### 4.C.2 Eye Tracking

We isolate head segment as matched by template. We upsample the head image, doubling image width and height, then we obtain a mean intensity value by sampling points along a elliptic arc passing through both eyes. We obtain 3 threshold values after ranking the sampled intensities the median, the 60 and the 85 percentile values, which are then used to to threshold the head image to segment the eyes at different intensities. The edges from each of the thresholded images is then extracted using Laplace edge detection. These are then combined and passed on to the ellipse detection algorithm. We isolate the detected edges from each eye separately by splitting the edges images into left and right panes. These are then independently processed by our customized implementation of a fast ellipsoid detection algorithm [58]. The ellipsoids are scored based on the number of edge pixels they overlap with, and the highest scoring ellipsoids are considered to provide the best fit for eye shape. The angle of each eye is then read out as the angle of each eye’s major axis relative to the body orientation. Noise from each of eye-angle trajectories is then suppressed during the data processing stage by passing the eye-data through a 4th order Butterworth low-pass digital filter (cut-off ≈ 28 Hz) Eye vergence is computed as left eye - right eye angle, letting clock-wise being positive angles.

#### 4.C.3 Tail motion

The tail motion is tracked by fitting a spine of 8 points that approximates tail length and curvature. We employ two methods for fitting the tail spine. The first is employed on every video frame and it uses pixel intensity to adjust spine points to detect tail midline by utilizing the fact that the tail appears brighter than background in our images. The image of the larva is Gaussian blurred, and the algorithm adjusts existing spine segment angles to track the center of mass of image intensity. Each spine segment is taken in order, starting from the most anterior-proximal spine segment, and then pixel intensity *i* is sampled along the arc of drawn from rotating the spine-segment across angles within *θ* = −40 ·… 40°from its current position. Centre of mass is calculated in vector containing 80 intensity sample points 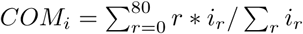 The segments orientation is fixed to the angle with the maximum sampled intensity. The second method is employed more irregularly (every 4 frames), and it aims to position the spine within the limits of the body contour but additionally to fix spine length appropriately. This method uses a variational method to best position the 8 point spine within a simplified, smoothed, larval contour shape made up of 90 points. By varying the spine parameters of length and angle, a Jacobian of matrix is computed and then a gradient in parameter space is computed. Gradient descent minimizes a cost function defined to be the sum of the distances of each spine point from the closest contour edge with an additional cost for fitting short tail lengths, therefore favouring the fitting of longer spines that are contained within the contour. The extracted angles of each spine segment is then noise filtered, during the data processing stage, through a Butterworth 4th order band-pass filter (4-123Hz).

#### 4.C.4 Detecting hunt events and labelling

Hunt events were detected via eye-vergence [6]. We define the start of hunting episode as the time when eyes verge beyond 45°with each eye being at least 19°inwards, and the end as the time when they diverge back out of the above range. Eye vergence needs to last at least 100 video frames (recording at 410 fps), and hunt-events need to be at least 300 frames apart, otherwise they are concatenated. The isolated video frames from the hunt events detected in evoked (prey) test conditions were played back to an independent observer, who was blind to the rearing group. They were allowed to observe the hunt-event for as long as they needed and then they were given a choice of labels to assign to the outcome of the event that included: *Capture success with strike, Capture success no strike, Capture failure no strike, Capture failure with strike, Failure no strike* (larva reached near the target and aborted or failed), *No target* (indicating events where no prey could be seen to be tracked).

### 4.D Bout detection and first turn to prey

Our data analysis was conducted via custom scripts in **R** [48]. We identify the start of hunting episode as the time when eyes verge beyond 45°with each eye being at least 19°inwards. Larval speed is measured by tracking the centre of the detected head template position (see above), and smoothed using a low pass Butterworth filter (24Hz). The capture bout is the last bout prior to the head of the larva passing from the position of the prey, and we exclude any bout that sometimes can occur immediately following prey capture [31]. In the case that hunting mode is initiated in conditions where there are multiple prey in the direction the larva faces we consider the hunt initiation to be prior to the turn that unambiguously identifies the prey item that is being tracked and which the larva will attempt to capture.

### 4.E Measuring larva size and prey distance

The distances are calculated based using an estimate of mm per pixel in the video calculated by measuring the diameter of the 35mm dish on screen in pixels. This mm/pixel ratio was estimated to be 35/790, i.e approximately 44*µm* per pixel, and this scaling was used for all measurements of length and distance from images. For reference, single pixel errors translate to an error distance of approximately 0.05mm.

Larval body lengths were measured on snapshots of larvae in straight posture by taking a straight line from the edge of the mouth-point to the point where the tail point vanishes Figure S13. Distances to prey were measured at the onset of the capture bout from the tip of the mouth point, while larval capture speed was tracked via a point on the head located near the anterior end of the swim bladder. Given our image resolution, single pixel errors are ≈ 10% when it comes to measuring distance from prey. Nevertheless, our measured distance from prey at the time of the capture strike are comparable to previously reported ranges [31].

### 4.F Bayesian statistics and notation

For a given dataset *D* described by a model *M*, Bayesian inference allows us to quantify the posterior distributions of model parameters (*θ*) according to the Bayes’ theorem

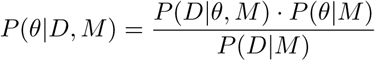

where the terms in the numerator correspond to the likelihood, *P* (*D*|*θ, M*), and the prior, *P* (*θ*|*M*), and the denominator is the normalization factor also known as model evidence.

Here we used the standard R package RJags [44] which allows us to estimate posterior distributions using the Markov Chain Monte Carlo method (MCMC). With this approach we can generate random samples distributed according to the posterior distribution of the model parameters and use them to calculate averages and uncertainties of relevant variables. In the next sections, when modelling prior distribution and data likelihood we used the following parameterization of standard probability distributions

- **Normal distribution**: 𝒩(*µ, t*), with mean *µ* and precision *t* = 1*/σ*^2^ defined as the inverse of the variance.
- **Negative binomial distribution**: NB(p, r), with 0 *< p* ≤ 1 being the probability, and *r* being the size parameter (*r* ≥ 0)
- **Gamma and inverse gamma distributions**: Gamma(*a, b*); InvGamma(*a, b*), with shape *a* and rate *b*.

### 4.G Statistical modelling of hunt rates

We counted the number of detected hunt-episodes recorded for each larvae in spontaneous and evoked test conditions (as described in Section 4.C.4) and then used Bayesian inference to estimate the mean hunt-event rate for each group. Here, a single Poisson is not sufficient to model the occurrence of hunt events in a group of larvae, but instead a model that assumes a mixture of Poisson processes of various rates *λ*_*i*_ is required to characterize hunt-activity in the group’s population. This mixture of rates aims to account for natural behavioural variability in hunt-rates, expected even among larvae of the same rearing group and test conditions. For this we employed a Gamma-Poisson mixture, where the number of hunt events *h* conditional to hunt rate *λ* in each recording time period (fixed to 10min) is distributed as a Poisson distribution

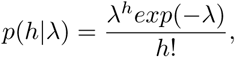

while hunt rates within the population are Gamma distributed

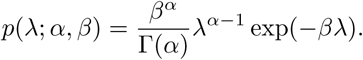

with hyperparameters *α* and *β*.

With this setting, the marginal distribution on the number of hunt events is a negative binomial NB(*r* = *α, p* = *β*(1 + *β*)^−1^), which provides a simple model with two parameters to characterize and regress each group’s mixture of hunt rates. In particular, we can express the mean hunt rate as

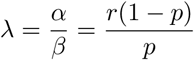

Once we infer appropriate parameters of NB(*r, p*) that fit the hunt rate data *h* of each group, we can then use those parameters to extract a distribution for the mean hunt-rate. Specifically, the model we used was *h* = NB(*r, q*) with priors for *q* = 𝒰(0, 1) and *r* = Gamma(1, 1), while the Gamma(*α, β*) distribution’s parameters, shape and rate respectively, can be recovered as *β* = *q/*(1 − *q*) and *α* = *r*. The model code is available on GitHub^1^.

### 4.H Statistical modelling of hunt duration

We model the total hunt frames that a larva spends hunting during the recording interval. We take the total number of frames in a video in which a larvae was engaged in hunting-mode to be a stochastic quantity composed of individual hunt-frame events that occur with a rate *λ*_*f*_. By counting the number of hunt-frames, the model of hunt-duration becomes equivalent to the hunt frequency model above, which counts the number of hunt-mode onsets for each larvae, and thus the negative binomial can be used here as well. Instead of counting hunt-events, we here count the total number of video frames each larvae spent in hunting mode. Note, we excluded larvae that produced no hunt events, and thus had a total hunt duration of zero frames. This is because hunt events are defined to have a minimum frame duration of 100 frames, and larvae without any detected hunt events make the hunt-duration distribution discontinuous. Using a model very similar to the one used for hunt-rates (Section 4.G) we estimated the density of mean hunt-duration through the negative binomial *µ* = *R*_fps_*r*(1 − *q*)*/*(*q*), where *R*_fps_ is the fps of video acquisition (410 fps), while *q* and *r* are the inferred model parameters from the duration data.

### 4.I Statistical modelling of hunt efficiency

We define the capture efficiency of each larva as the fraction of hunt events it performed that ended with successful capture, over the total number of hunt events in which prey capture was attempted. Larvae which did not perform any capture attempts are excluded from this analysis.

We utilized the earlier hunt-rate model (Section 4.G), to model the distribution of capture attempts *h* of each larval group,

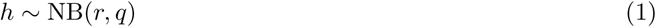

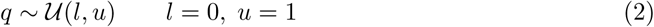

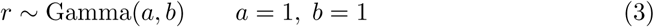

here it is extended to infer the probability *p*_*s*_ of successfully capturing a prey. For this, we used a binomial distribution to model the number *N*_*s*_ of successful captures in the 10 min recording period

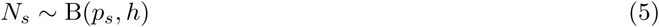

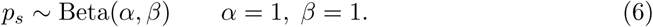

which depends on the total number of events *h*. By using Bayesian inference we estimated the distribution of capture probability *p*_*s*_ for each group independently.

### 4.J Linear regression of turn to prey behaviour

The data used for this model are the magnitude of the first turn towards prey *ϕ* and the prey azimuth *θ*, prior to the larva turning towards the prey. Data points are pooled for each rearing group. We characterized each group independently using a linear model and performed a Bayesian analysis of the linear coefficients, which enables us to compare rearing groups using their posterior distributions. The linear regression model for each group is defined as:

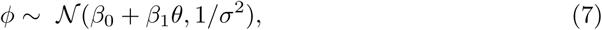

which assumes the errors are independent and identically distributed as normal random variables with mean zero and variance *σ*^2^, for which we used an inverse gamma prior

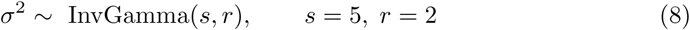

while for the linear coefficients we used

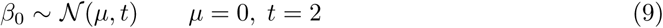

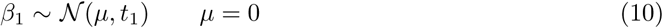

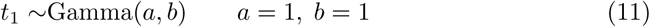

Our aim is to update the initial guess distributions for the model’s *β*_0_, *β*_1_, *σ*^2^ for each rearing group according to its relevant data points. We then report and compare the density estimates for the slope parameter *β*_1_ of each group’s model, after smoothing using a Gaussian kernel (BW= 0.01), on ***Figure 5C***.

### 4.K Classification of fast and slow capture swims

The final motion with which a prey is captured is executed with a range of speeds indicating difference in vigour. Broadly speaking we observe two types of capture motions, a stereotyped fast speed capture swim, which is usually successful executed with the larva standing at some distance to the prey, and a weak bite capture, which can only be successful if executed when the prey is very close or touching the larva.

Our tracking system is not able to automatically give us reliable and accurate information on the distance to prey prior to capture, as small position errors get magnified due to low video resolution. Thus we decided to conduct supervised tracking of the specific hunt events, to minimize errors but also verify the validity of our data. The video frames during final capture were re-analysed and the distance from the edge of the mouth-point *d* to the prey position was measured with the help of a user distance measurement tool that we built into our tracker (see Section 4.E). These distances were then used as data on distance to the prey at prior to the capture bout. Capture speed *s* is measured as the peak speed of the final bout occurring between the frames starting from the frame of capture bout onset (on which the prey-distance is manually measured) and up to the first time the larva’s speed goes below a motion detection threshold (4mm/sec) in the frames proceeding the time of when the larva’s centroid has reached the closest point to the prey.

Capture speed *s*^*c*^ and distance to prey *d*^*c*^ were modeled using a mixture of two bi-variate normal distributions to accommodate slow and fast events labeled using the index *c* = {*s, f* }

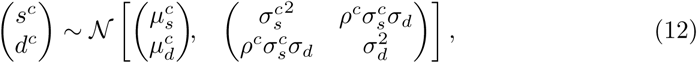

where 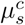 and 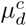 are respectively the mean speed and distance to prey, 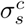 and 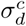 the standard deviations and *ρ*^*c*^ is the correlation coefficient between speed and distance for each cluster *c*. We used the following priors on the mean and covariance parameters

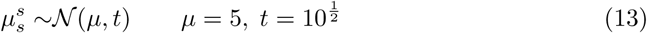

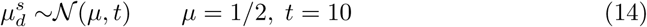

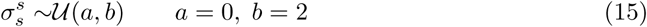

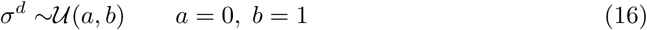

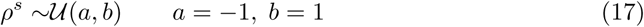

for the slow group and

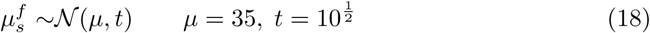

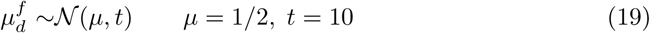

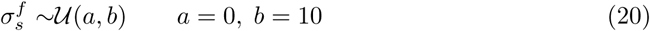

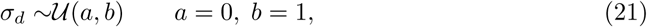

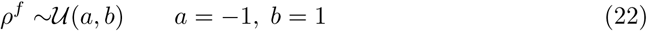

for the fast group, with a narrow *σ*^*s*^ to add the prior belief that low capture speeds are positioned in narrow range at the lower end in the range of capture speeds observed. The prior on group membership was 0.5 for fast and slow capture.

The standard deviation for capture speed is wider and is representing the prior belief that fast capture speeds occupy a wide range of speeds above the slow captures modes. Indeed beyond slow captures, capture speeds do not appear stereotyped and are seen to vary probably in relation to other parameters such as distance to prey. Note the prior for distance to prey is set identically for both clusters and as such we expect the data to inform the mean distance of each cluster.

The probability of membership on either cluster *c* ∈ 0, 1 for fast/slow is estimated from a normal distribution with as *p*_*f*_ = 𝒩 (∑ *I*(*c*_*i*_ = 1)*/N*, 0.03), where *I*(*c*_*i*_ = 1) is equal to 1 if data point *i* has been assigned to the fast cluster. Data points were assigned to the fast cluster if the expected cluster membership label was *E*[*c*] *>* 0.7, otherwise they were considered as slow capture swims.

### 4.L Statistical modelling of group behaviour

We built a hierarchical statistical model to estimate mean group behaviour that is based on model estimates of mean hunting behaviour per larva. This is a generalization of our earlier model of capture-speed and distance, only here the top level structure is single multivariate normal distribution that models *X*^*g*^, a vector containing estimates of mean capture speed *S*, distance to prey *D* and turn-ratio *T* for each larva. Details of this model and example code can be found in the code repository ^2^.

### 4.M PCA of larval hunting behaviour and predicting efficiency

Principal component analysis can reduce the dimensions needed to describe a large set of correlated predictor variables to a smaller, less correlated set of covariates, that nevertheless maintains most of the information in the larger set. A subset of the resulting covariate components can then be used to regress an outcome variable, effectively producing a model that predicts a response based on a subset of principal components.

The process of obtaining principal components involves constructing a covariance matrix **A** of our observation data and then calculating its eigen decomposition. For our analysis we constructed a matrix of vectors, each one representing the hunt behaviour of each larvae estimated from measurements taken from successful hunt episodes alone. For each larva *i* we defined vector 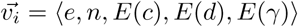, where *n* = *N*_*s*_ + *N*_*f*_ is the capture attempts recorded for larva *i, e* = *N*_*s*_*/n* denotes its capture efficiency, *E*(*d*) is the mean distance to prey estimated from the onset of the capture bout across a larva’s hunt events, and *E*(*γ*) is mean turn-ratio at the initial turn towards prey. For ***Figure 7B*** we derive the expected values per larva from the results of our earlier group model (see Section 4.L), while in ***Figure S9*** we used empirical means (*E*(*x*) = ∑_*n*_ *x*_*i*_*/n*) to estimate the mean hunt behaviour. In order to ignore any scale effects of covariance on PCA, we standardized the variables using

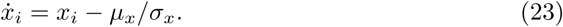

With the exception of turn-ratio where *µ*_*x*_ = 1, *µ*_*x*_ was set to the mean value of each behavioural variable *x* calculated across larval vectors.

Each principal component packs correlated variables that could possibly act as better predictors and provide compact regression models in situations where there many predictor variables and relatively few samples. We used the *pls* package [54] in **R** [48], to conduct principal components regression of larval efficiency using a linear model and the principal components of matrix of hunt behaviour estimators of 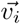. The algorithm reported that using 3 PCs gave the smallest root mean square prediction error with coefficient of variation CV=0.1335, capturing 31.7% of efficiency variance.

## 5 Acknowledgments

We’d like to thank Sabine Issop for help with manual labelling of hunt events, the fish facility staff at King’s College London for their excellent fish husbandry, Elina Jacobs and Adil Khan of King’s College London for their feedback on our manuscript.

This work was supported by a Wellcome Investigator Award MPM: 204788/Z/16/Z)

## 6 Supporting Information

**Figure S1.**
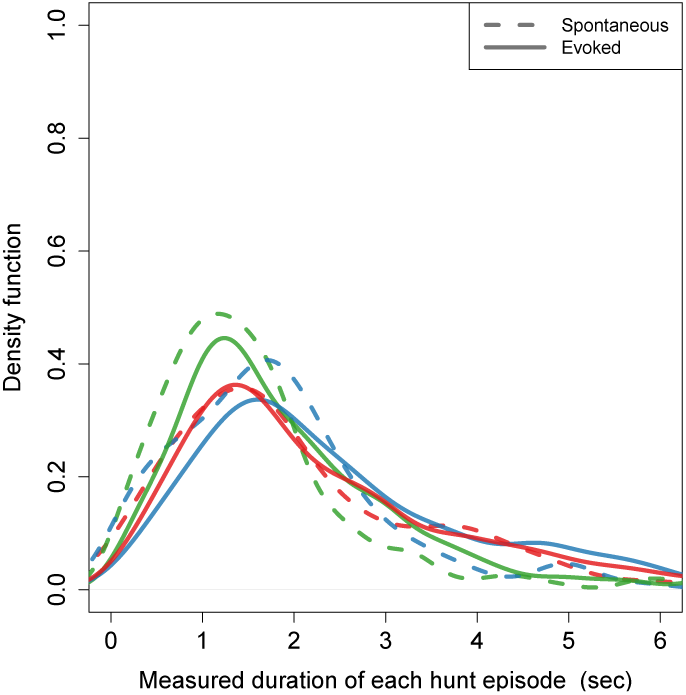
The duration of individual hunting episodes are similar between the spontaneous and evoked events of each group. Raw data kernel density (BW=0.25 sec) estimation of, from *n*_*s*_ = 699 spontaneous (*n*_LF_ = 234, *n*_DF_ = 227, *n*_NF_ = 238) and *n*_*e*_ = 2578 evoked (*n*_LF_ = 941, *n*_DF_ = 797, *n*_NF_ = 840) hunting events. Distributions across conditions and groups show large overlap in episode duration. The LF group displays somewhat shorter mean episode duration in spontaneous conditions (mean duration LF 1.74 ± 0.08 sec, DF 2.04 ± 0.08 sec., NF 2.27 ± 0.11 sec.). 95% of the spontaneous hunt-episode duration data are below 3.9 sec. for LF, while this is 92% of DF, and 84% for NF. The NF group’s 95% percentile is at 5.02 sec. indicating that there are some comparatively longer spontaneous events in this group. In evoked conditions, the average duration of each hunt-event in LF group’s is shorter (LF 2.07 ± 0.04 sec., DF 2.62 ± 0.06 sec., NF 2.51 ± 0.06 sec.), with 95% of data below 4.6 sec. for LF and DF. Similar to spontaneous conditions, the NFs 95% percentile sits higher, at 5.8 sec, indicating again that NF hunt-events can be rather sustained compared to LF and DF.

**Figure S2.**
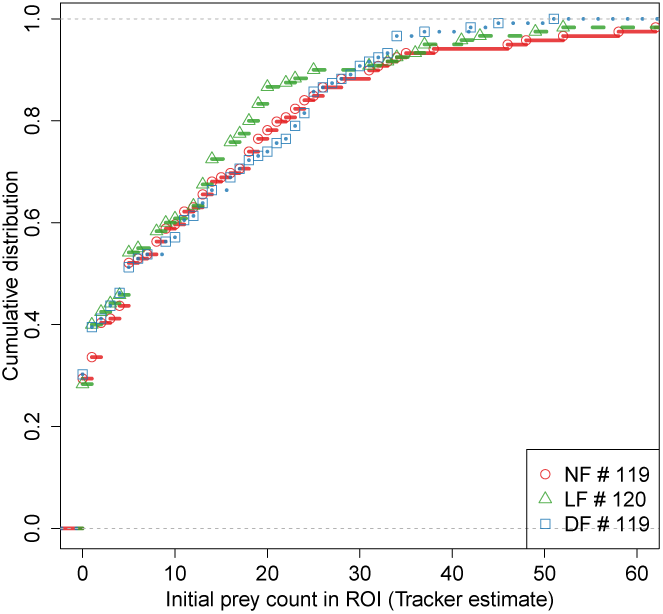
CDF of number of detected prey at the beginning of each experiment, shows that the evoked hunt-rates were tested in similar prey density conditions between groups.

**Figure S3.**
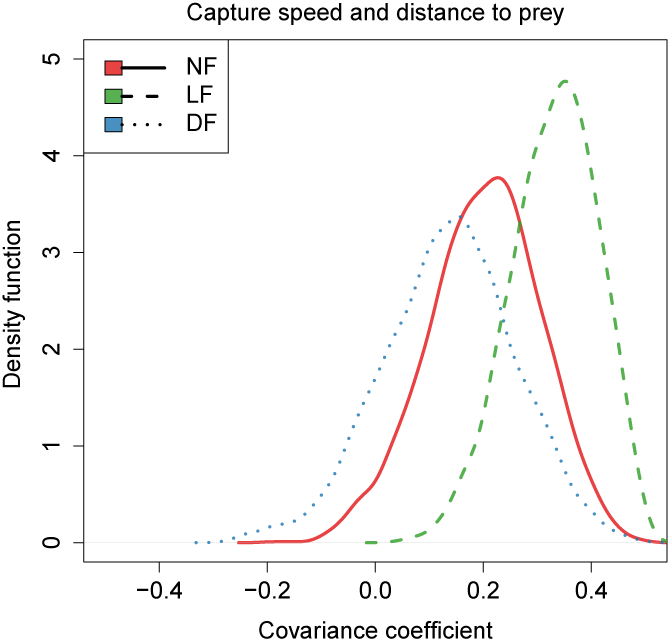
A relationship between distance and speed is also captured in the covariance of the fast-cluster’s model.

**Figure S4.**
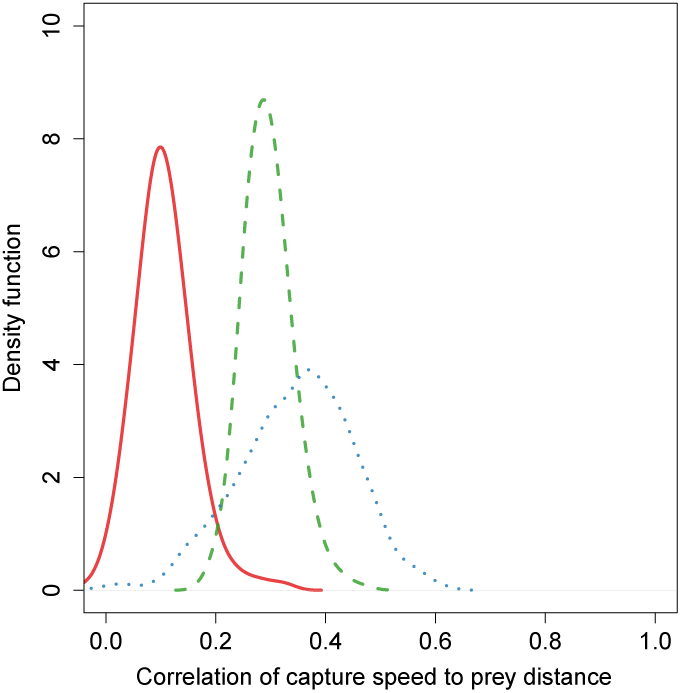
Bootstrapping the speed-distance correlation in the data points of the fast-cluster alone also reveals correlation.

**Figure S5.**
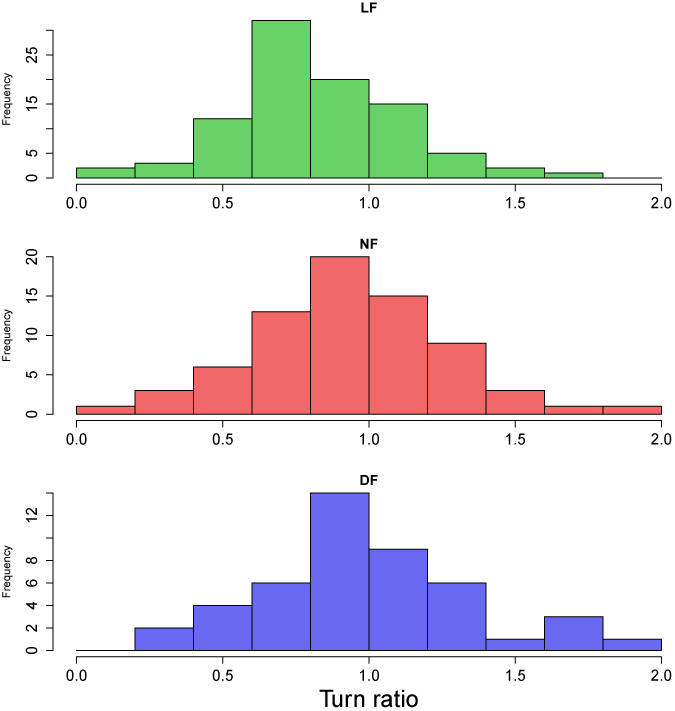
Histogram of turn ratio data also shows that hunt events from the LF group have a bias towards turn ratios below unity (undershooting).

**Figure S6.**
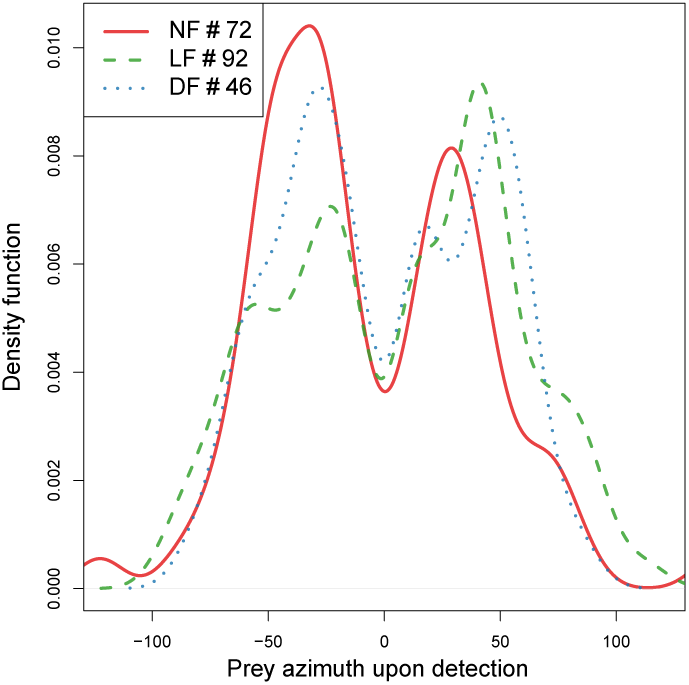
Prey azimuth at start of successful hunting event. Differences in under-shooting are not due to differences over the angle at which prey are being detected. Estimated density of prey detection angle (Bw=10°) shows that the hunt data across groups does not differ in the angle of initial prey detection, having similar bimodal distributions with peaks around 30°-50°in azimuth.

**Figure S7.**
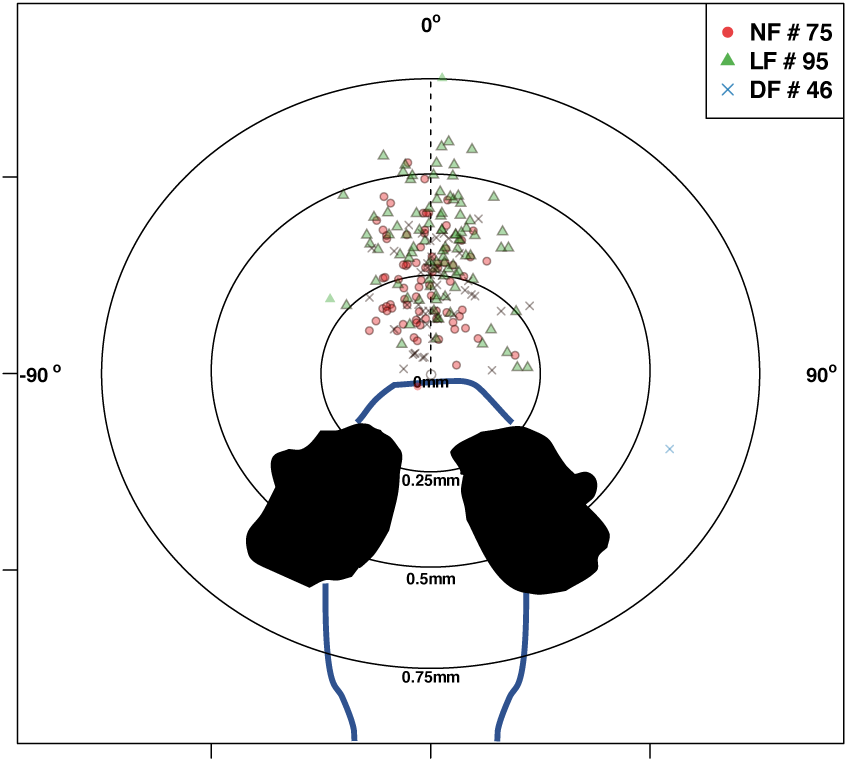
Differences in turn ratio do not alter the positioning of prey prior to successful captures. These are similarly located within the anterior field zone of eye vergence across hunt events from all groups, yet at different distance from the mouth (see Figure 4F).

**Figure S8.**
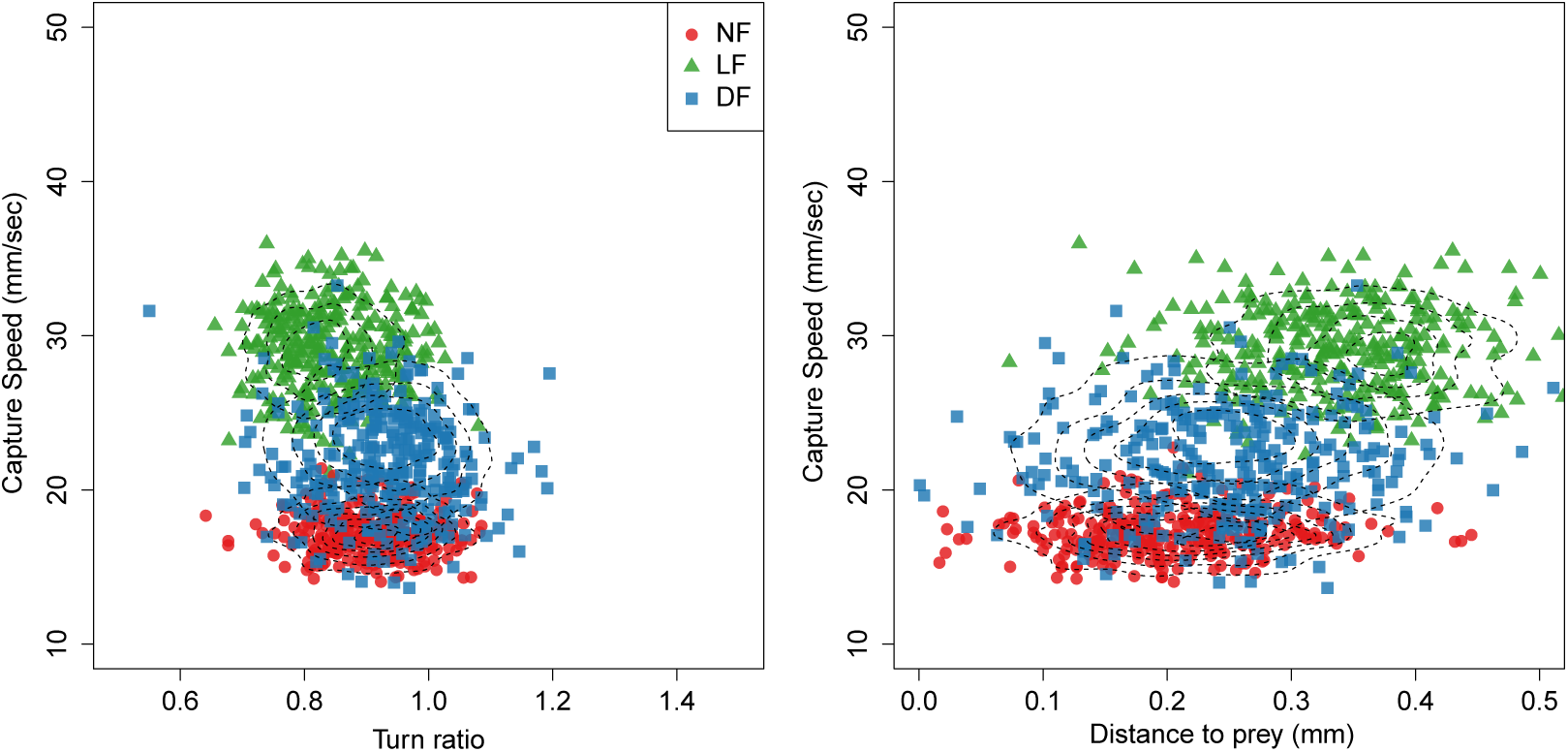
Detailed 2D views of ***Figure 7A*** that compare the parameter distributions between models of group hunt behaviour. The LF group has a distinct behavioural signature compared to controls verifying that experience modifies larval behaviour. Also, larval behaviour is more consistent within the LF group compared to the wider distributions of uncertainty over DF and NF behaviour.

**Figure S9.**
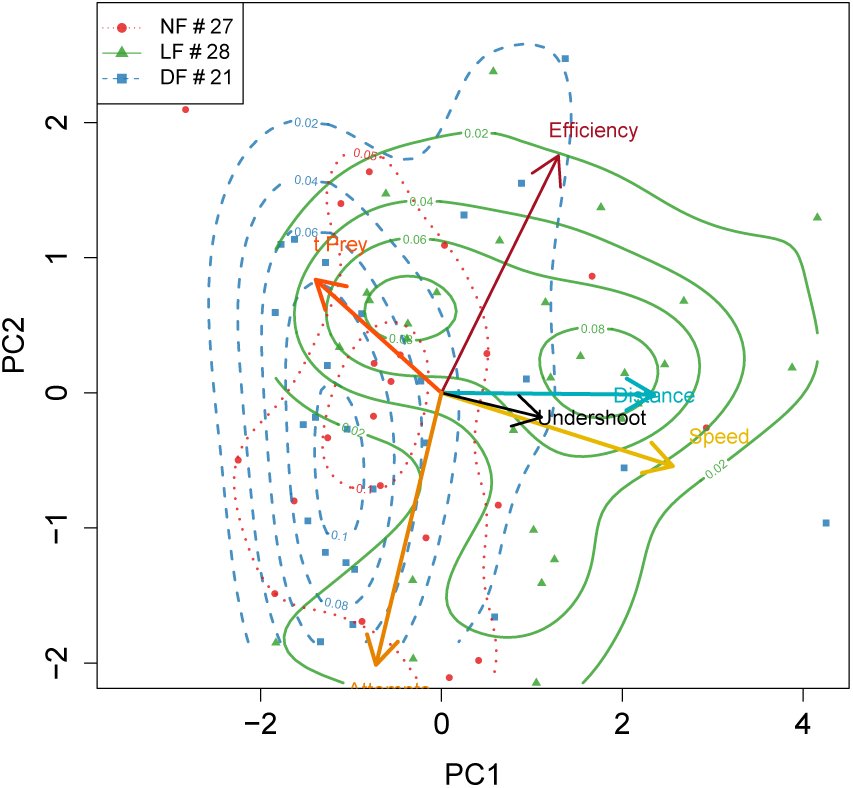
PCA applied to mean larva behaviour calculated empirically showing general agreement with model based in ***Figure 7B***

**Figure S10.**
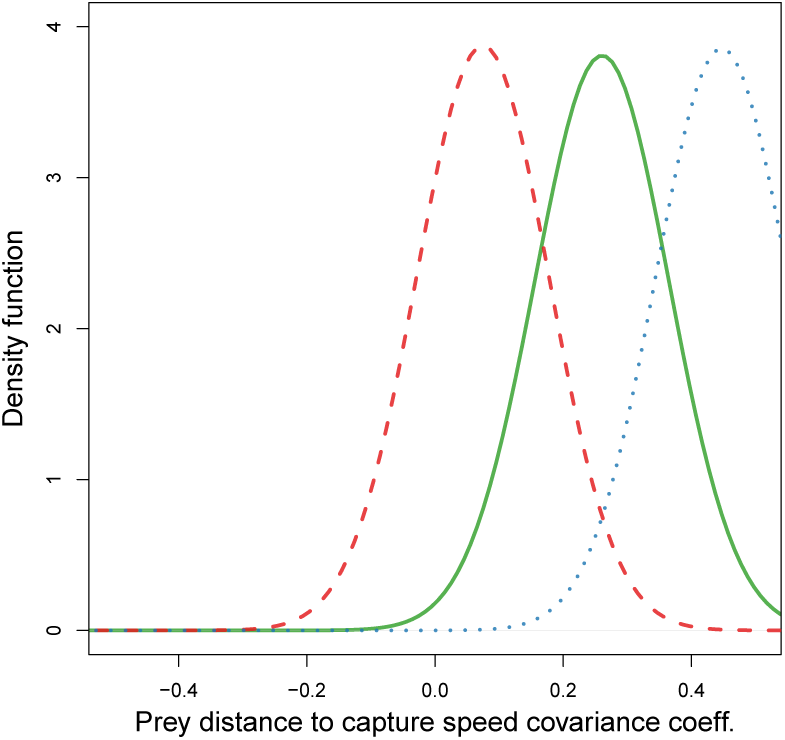
Covariance describing the relationship between capture speed and prey-distance provides evidence that larval groups differ in how they adjust capture swim speed against distance to prey. The model provides stronger evidence for a relationship in the DF and LF group, while NF larvae may be frequently choosing capture speed without reference to prey distance.

**Figure S11.**
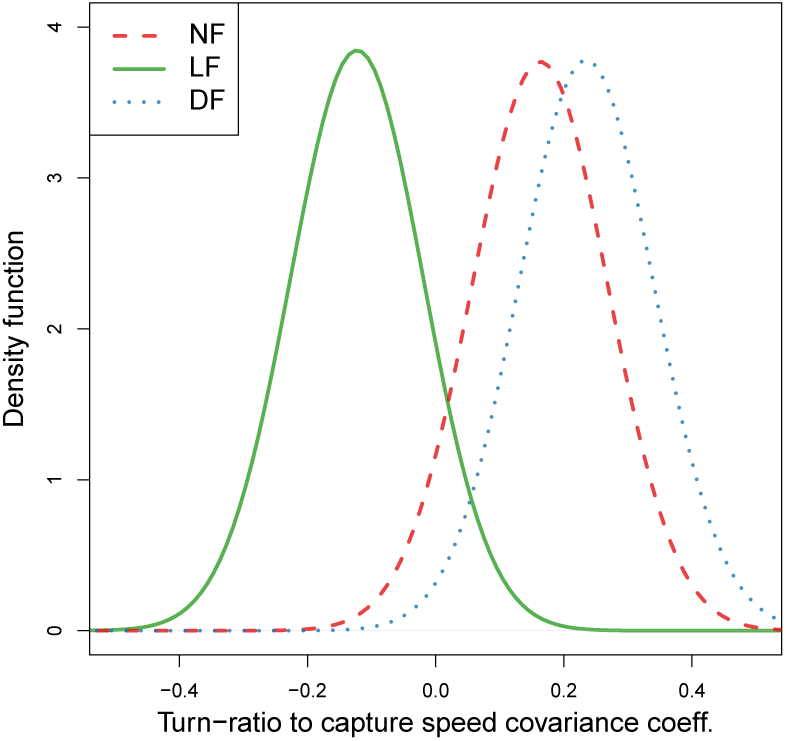
Model covariances confirm that capture speed and undershoot are correlated in experienced larvae (LF). In contrast, the opposite tendency, to overshoot, is associated with fast capture swims in control group larvae.

**Figure S12.**
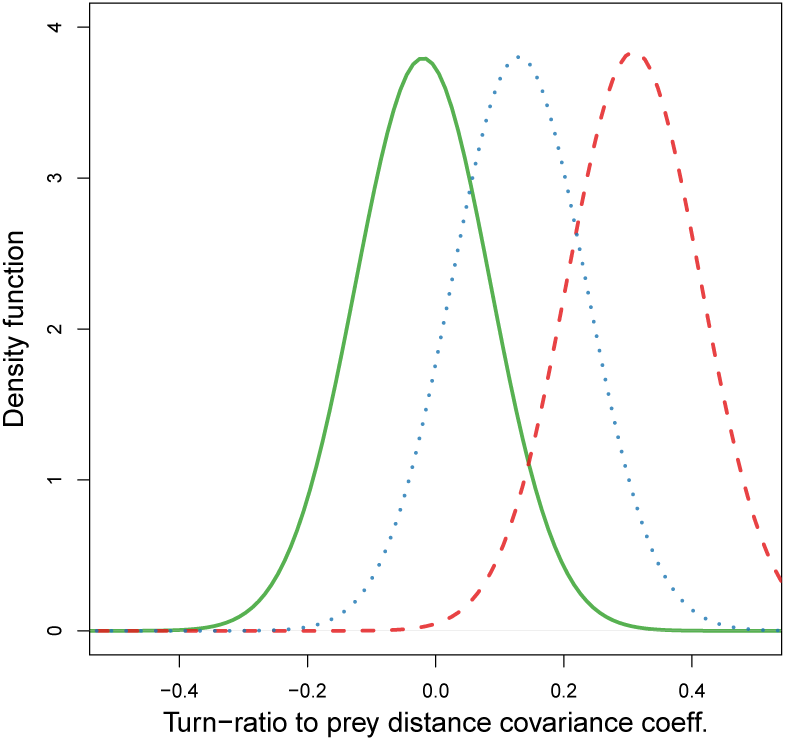
Initial turn-to-prey and capture distance are mostly unrelated for LF and DF, yet evidence that when NF fish overshoot they end up at longer distance to prey exists.

**Figure S13.**
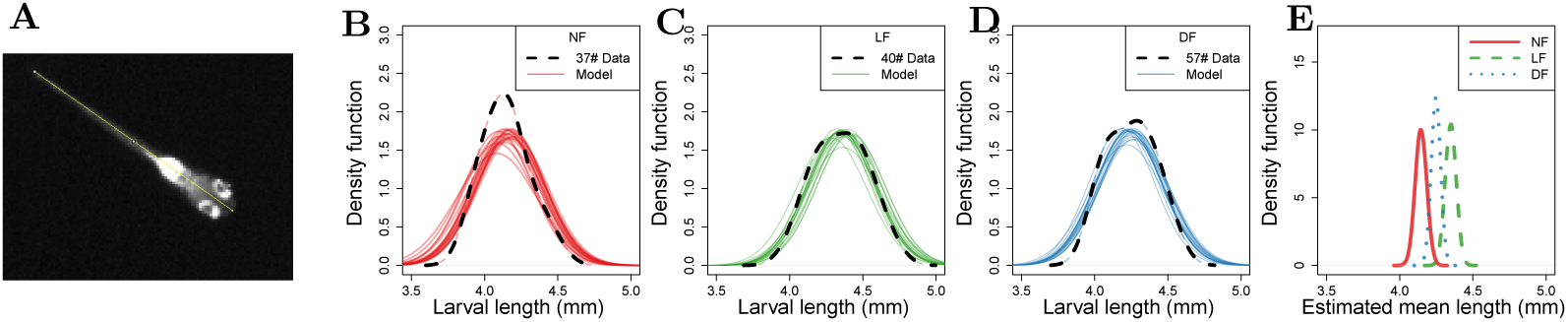
Analysis of larval length shows small statistical differences in mean length, with larvae on live-prey being the longest. **(A)** Larval standard length is measured in pixels from mouth point to edge of tail on video frames when the larva is not in hunting mode and in straight posture. This is then converted to mm by using an estimate of mm/px. **(B-D)** Distribution of body lengths in the different feeding regimes. Dotted black lines indicate kernel density smoothed distributions of measured larva lengths (Gaussian kernel BW=0.1) and solid lines show 30 samples of likely body length distributions based on a Gaussian model fit, whose parameters (*µ,σ*) were estimated from the data using Bayesian inference. **(E)** The estimated probability densities of mean body length. The probability density functions for each feeding group are distinct suggesting small differences in mean body length.

**Figure S14.**
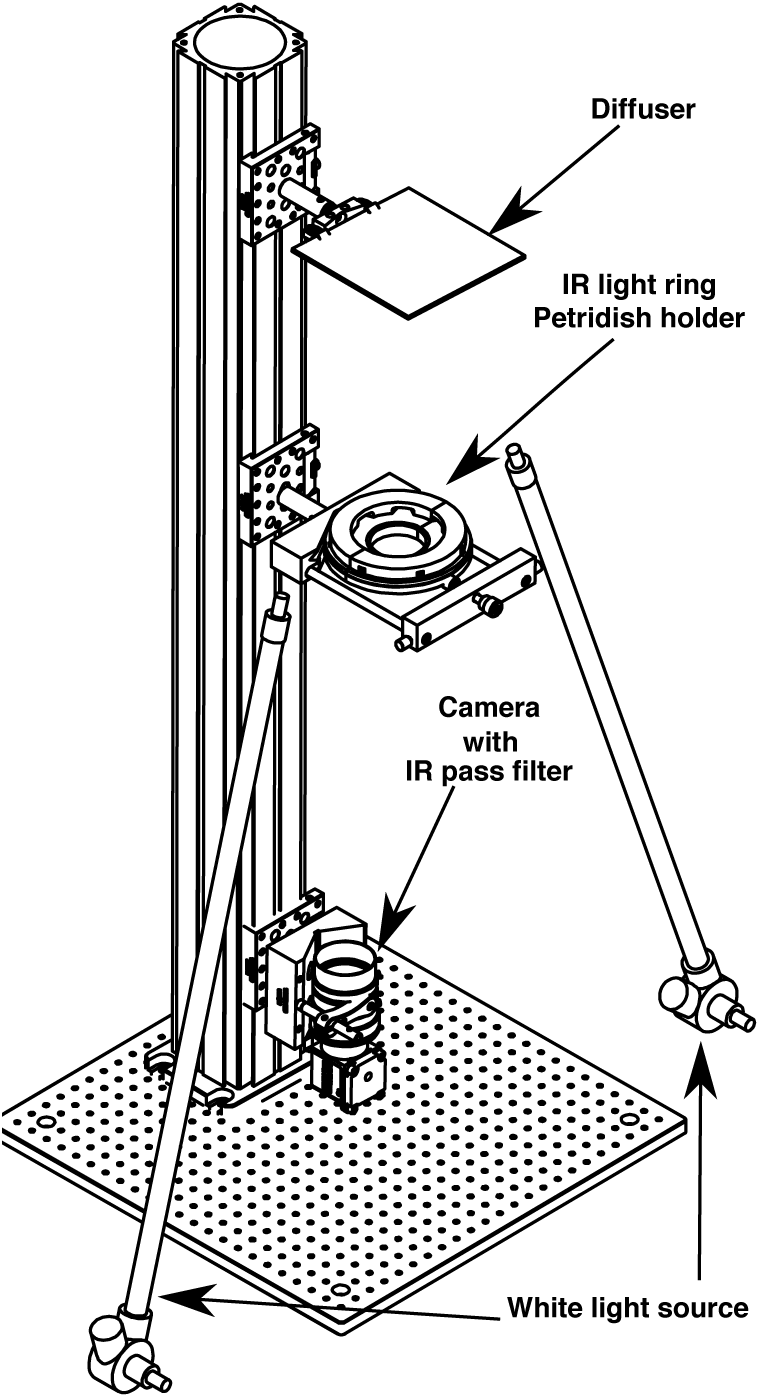
Behaviour imaging system and example images of hunting sequence from tracking software. The behavioural imaging system is connected to recording software that initiates recording when the fish is within circular region of interest (ROI), set such as behaviour is recorded only when larval is sufficiently away from the edge of the petridish. The recording session timeout is 10min, beyond which time new recording events are not triggered, The maximum duration of recording events is limited to 2 mins, beyond which time if the larva is still in the ROI and within the 10min timeout a new event is recording is triggered.

**Figure S15.**
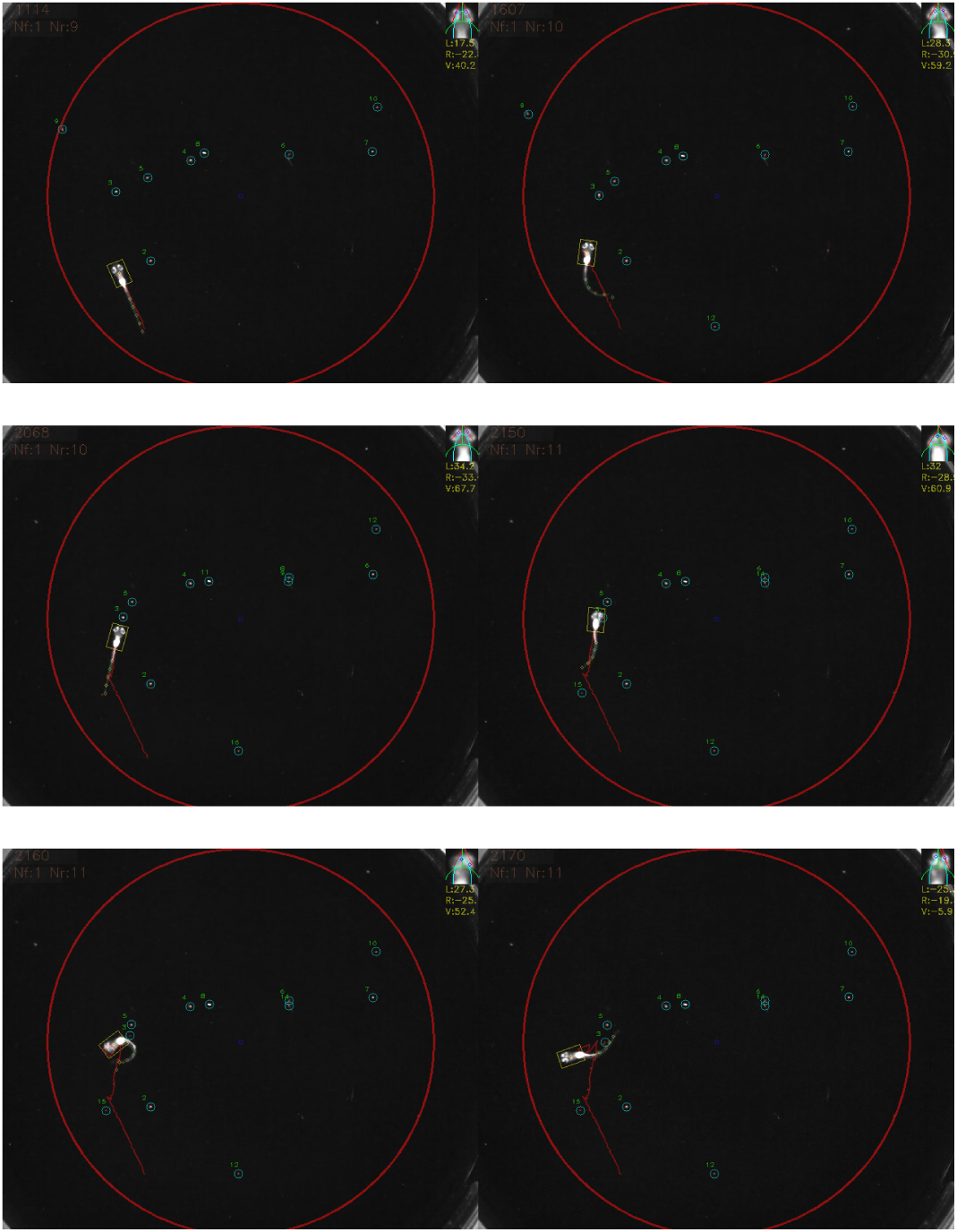
Example frames from a hunting sequence being tracked via our software showing initial detection of prey, turn to prey, approach and capture. The eye vergence angle is detected and shown at the top right of the screen.

**Figure S16.**
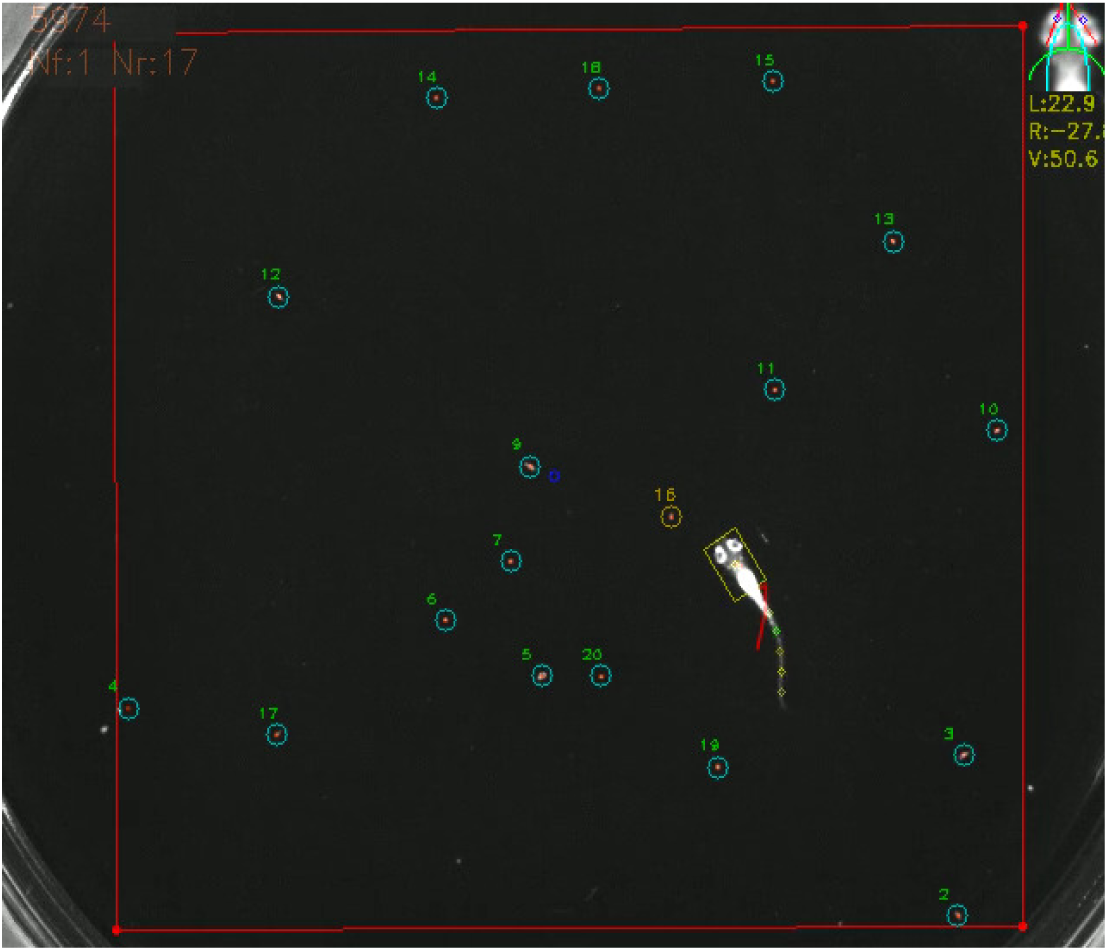
Example video of a successful hunt episode being retracked along with the target prey.

**Figure S17.**
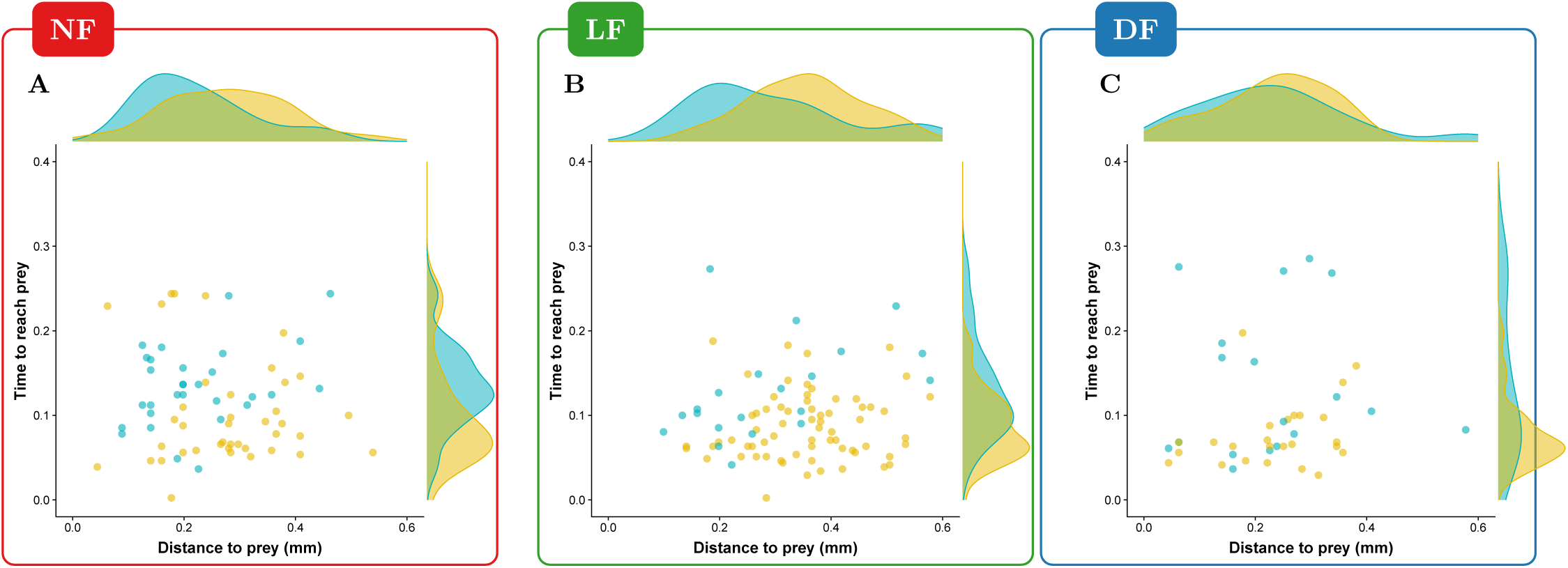
The time it takes to reach prey for successful fast capture swims does not strongly depend on the distance from which these are executed. Points are coloured according to the classification in capture speed as in Figure 4 Yellow: fast-capture swims, cyan:slow capture swims. In all groups the time to reach prey is longer for the capture swims that were clustered as slow, while for fast-captures swims the timing is more compactly distributed below 0.15 sec. Maintaining such timing would require adjusting capture speed with prey-distance. It appears that LF **B** can regularly do this even for prey distances beyond 0.4mm, where successful hunt episodes from NF, DF (**A,C**) are rare.

## Footnotes

1 https://github.com/kostasl/ontogenyofhunting_pub.git

2 https://github.com/kostasl/ontogenyofhunting_pub/blob/master/stat_3DLarvaGroupBehaviour.r

